# Collicular circuits for flexible sensorimotor routing

**DOI:** 10.1101/245613

**Authors:** Chunyu A. Duan, Marino Pagan, Alex T. Piet, Charles D. Kopec, Athena Akrami, Alexander J. Riordan, Jeffrey C. Erlich, Carlos D. Brody

## Abstract

Historically, cognitive processing has been thought to rely on cortical areas such as prefrontal cortex (PFC), with the outputs of these areas modulating activity in lower, putatively simpler spatiomotor regions, such as the midbrain superior colliculus (SC). Using a rat task in which subjects switch rapidly between task contexts that demand changes in sensorimotor mappings, we report a surprising role for the SC in non-spatial cognitive processes. Before spatial response choices could be formed, neurons in SC encoded task context more strongly than neurons in PFC, and bilateral SC silencing impaired behavioral performance. Once spatial choices could begin to be formed, SC neurons encoded the choice faster than PFC, while bilateral SC silencing no longer impaired choices. A set of dynamical models of the SC replicates our findings. Our results challenge cortically-focused views of cognition, and suggest that ostensibly spatiomotor structures can play central roles in non-spatiomotor cognitive processes.

## INTRODUCTION

Our response to the sensation of our phone ringing will be very different in the context of having just heard “the play is beginning, please silence all phones” versus a context in which we have just been told “your child’s school is about to call with urgent information.” Such top-down flexible switching in sensorimotor routing is central to our daily lives, and is a core component of what is known as executive function (Miller and Cohen, 2001). Motivated by seminal studies where patients with prefrontal cortex (PFC) lesions (Guitton et al., 1985; Pierrot-Deseilligny et al., 1991a) or schizophrenia (Fukushima et al., 1988; Hutton and Ettinger, 2006) failed to perform flexible task switching, decades of primate research has focused on the PFC as the pivotal neural substrate underlying executive functions, especially the inhibitory control of context-inappropriate behaviors (Lo and Wang, 2016; Munoz and Everling, 2004; Pierrot-Deseilligny et al., 2003). Downstream subcortical regions, such as the midbrain superior colliculus (SC), were hypothesized to instantiate reflexive sensorimotor pathways that, depending on context, may need to be suppressed by cortical input (Everling et al., 1998). A small number of studies showed data inconsistent with the cortical inhibition model (Condy et al., 2007; Wegener et al., 2008) or even demonstrated an inhibitory role of the human SC (Pierrot-Deseilligny et al., 1991b). However, until recently (Everling and Johnston, 2013; Johnston et al., 2014), these results remained largely overlooked and the prefrontal inhibition model continued to be the dominant hypothesis for how executive control is implemented in the brain. Furthermore, none of these studies provided high temporal resolution perturbation data in the SC, therefore lacking direct evidence for an active role of SC in specific, differently timed, processes of executive control.

The spatiomotor view of the SC is consistent with traditional views of brain function that separate cognitive operations from downstream motor implementations (Miller and Cohen, 2001; Pierrot-Deseilligny et al., 1991a; Sigman and Dehaene, 2005), as well as with a large body of work in the SC centered on its spatiomotor functions (Gandhi and Katnani, 2011). Classic studies of the SC intermediate and deep layers have focused on their critical role in generating spatial orienting movements in learned behaviors (Felsen and Mainen, 2008; Robinson, 1972; Schiller and Stryker, 1972; Sparks and Hartwich-Young, 1989; Wurtz and Goldberg, 1972), as well as in innate behaviors (Dean et al., 1989; Evans et al., 2019). Recently, a more integrative role of the SC in spatial target selection, spatial attention and spatial decision-making has been proposed (Basso and May, 2017; Krauzlis et al., 2013; Wolf et al., 2015), but all strictly within the confines of spatial processing.

Here, we report evidence that the SC encodes, and is required for, non-spatiomotor processes. We recorded neural activity of SC and PFC populations in rats performing a rapid task switching behavior (Duan et al., 2015), where rats exert executive control and demonstrate behavioral hallmarks that closely parallel those found in analogous primate paradigms (Everling and DeSouza, 2005; Munoz and Everling, 2004; Weiler and Heath, 2012). In addition, taking advantage of high time resolution optogenetic silencing tools available in rodents, we examined the requirement of the SC for different phases of the task. We found that during a non-spatial task encoding delay period, before spatial response choices could be formed, the SC population contained significantly more information about non-spatial task context identity than the PFC population. Moreover, bilateral silencing of the SC during the same non-spatial delay period impaired behavioral accuracy. In contrast, bilateral silencing that began once spatial choices could begin to be formed did not impair choice accuracy. This was true even though during this period SC neurons contained earlier and stronger information than PFC neurons about the upcoming spatial response choice, which suggested that the SC plays an important role in generating those spatial choices. To account for these behavioral, electrophysiological, and perturbation data we developed a dynamical model framework and found numerous, highly varied model circuit architectures that were compatible with these results, with the SC models involved in both generating the spatial choices and in non-spatial task context processing. Despite heterogenous parameter values and circuit dynamics, the many model solutions consistently performed the behavior by encoding task context and inhibiting task inappropriate responses within the SC itself, in a manner consistent with our electrophysiological and optogenetic data. These results argue against cortical inhibition of collicular activity as the key mechanism for executive control (Pierrot-Deseilligny et al., 1991a; Ridderinkhof et al., 2004), and instead suggest that the active representation of the non-spatial, relevant task goal in SC neurons is essential for flexible sensorimotor routing.

## RESULTS

### SC and PFC populations encode task context and motor choice during rapid task switching

We trained rats to perform rapid task switching in a behavior in which two different task contexts require opposite sensorimotor orienting responses (Duan et al., 2015). On each trial, rats are first presented with an non-spatial auditory cue indicating the task context in effect for the current trial (labelled either the ‘Pro’ or ‘Anti’ context). This is followed by a silent memory delay period, and then by a spatial choice period during which a visual stimulus to one side is turned on; in the Pro context rats are required to orient toward it while in the Anti context they should orient away (Figures 1A and 1B). This final choice period, beginning when the side light is turned on, is when spatial sensory information first becomes available, to be combined with the non-spatial task context information in order to perform the correct sensorimotor transformation and produce the correct spatial orienting response. We previously found that rats can flexibly switch between these two task contexts from one trial to the next, and display multiple behavioral asymmetries between Pro and Anti responses (Figure S1; Duan et al., 2015), that are similar to those observed in a closely related primate paradigm (Everling and DeSouza, 2005; Munoz and Everling, 2004; Weiler and Heath, 2012). These asymmetries indicate the Pro task as a stimulus-driven task, while the Anti task is more cognitively demanding (Duan et al., 2015).

**Figure 1.**
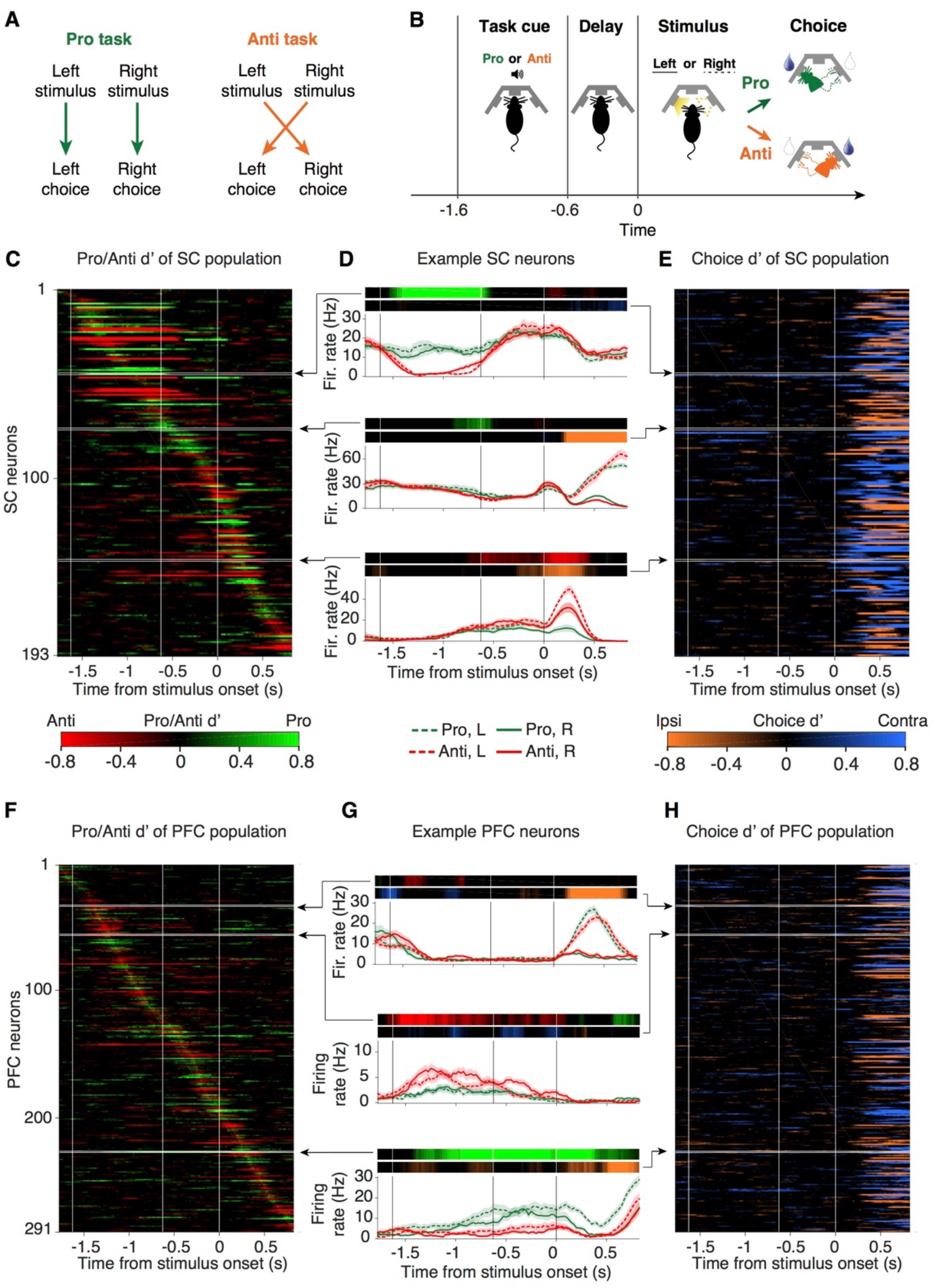
Individual SC and PFC neurons encode task and choice variables during flexible sensorimotor routing. (A) Rules for the Pro and Anti task contexts. In the Pro task, rats should orient *toward* a lateralized stimulus (left or right) for reward; in the Anti task, rats should orient *away* from the stimulus for reward. Trained rats can switch between these two known task contexts from one trial to the next. (B) Rats nose poke in the center port to initiate each trial and keep fixation during the task cue (Pro or Anti sound) and delay periods. After the delay, the animal is allowed to withdraw from the center port, and a lateralized light (left or right) is turned on to indicate the stimulus location. Rats then poke into one of the side pokes for reward. (C) Matrix of Pro/Anti selectivity for the SC population. Each row of the matrix represents the Pro/Anti signed d’ of a single neuron as a function of time. Neurons are sorted by the timing of their peak Pro/Anti absolute d’. (D) Peri-stimulus time histogram (PSTH) for 3 example SC neurons on Pro-Go-Right (green solid), Pro-Go-Left (green dashed), Anti-Go-Right (red solid), and Anti-Go-Left (red dashed) trials. PSTHs are aligned to stimulus onset. Top panel, Pro/Anti signed d’ and Go Left/Right signed d’ as a function of time for each neuron. (E) Matrix of Choice (Go Left/ Go Right) selectivity for the SC population. The recorded hemisphere was randomly assigned for each rat. We therefore quantified choice selectivity as preference for orienting responses ipsi- or contralateral to the recorded side. Neurons are sorted as in panel C. (F-H) Same as in (C)-(E), for the PFC population.

To investigate neural representations in SC and PFC, we recorded well-isolated single units in the intermediate and deep layers of the SC (215 neurons; Figure S2; STAR Methods), and in the prelimbic region of the medial PFC (331 neurons), from 7 rats performing the ProAnti task-switching behavior. Individual neurons in the SC and PFC displayed firing rates that depended on, and therefore encoded, task context (Pro or Anti) as well as the subsequent spatial motor choice (Left or Right; Figures 1C-H). Similar to observations in prefrontal cortex during cognitive tasks (Figures 1F-H; Durstewitz et al., 2010; Johnston et al., 2009; Karlsson et al., 2012; Mante et al., 2013; Rigotti et al., 2013; Rodgers and DeWeese, 2014; Schmitt et al., 2017), the encoding in SC appeared to be multiplexed and highly heterogeneous across different neurons (Figures 1C-E; Chan et al., 2017; Everling et al., 1999; Felsen and Mainen, 2012). In both populations, we observed neurons that fired more on the Pro or the Anti task trials (Figures 1C and 1F); and neurons that were selective for orienting responses ipsi- or contralateral to the recorded side (Figures 1E and 1H; Erlich et al., 2011; Felsen and Mainen, 2008). In contrast to past rodent studies of the SC using delayed response tasks, the encoding of ProAnti task context here is dissociated from motor planning, as supported by the lack of spatial coding before the choice period (Figures 1E and 1H). An initial inspection of the entire SC population revealed no apparent correlation between task and choice preference (Figure S4A) or clear temporal order of Pro versus Anti task selectivity. Given the heterogeneous and multiplexed nature of single neuron responses (Figures 2A and 2B), we focused on the representation of task variables on the population level (Figure 2).

**Figure 2.**
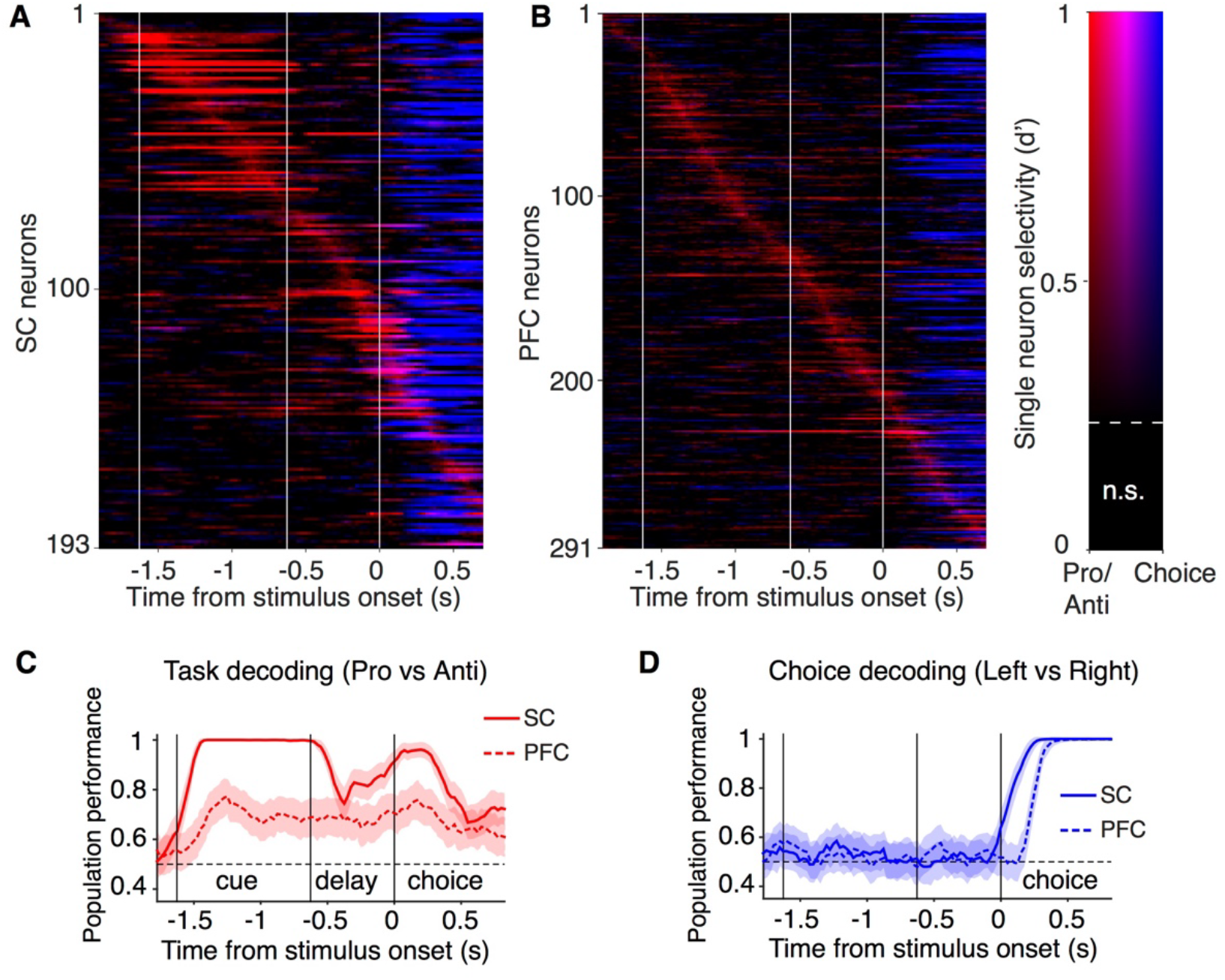
SC population contains stronger task context information and earlier choice information than PFC population. (A-B) Multiplexed task and choice encoding in the SC (A) and PFC (B) populations. Similar to Figures 1C-H, but instead of focusing on the sign of task and choice preference, the strength of task and choice selectivity are simultaneously illustrated here for each neuron to demonstrate multiplexing. The intensity of the color represents how “informative” a neuron is, and the RGB values are associated with different types of information (Pro/Anti, red; choice, blue; mixed, purple). d’ that are not significant (n.s.) are set to 0. (C-D) Evolution of classification performance over time in the SC (solid) and PFC (dashed) population. C, mean ± s.d. performance for linear classification of correct Pro versus Anti trials. Spikes are aligned to stimulus onset, and counted over windows of 250 ms with 25-ms shifts between neighboring windows. Note that performance is plotted over the right edge of the window (causal). D, classification performance to linearly separate Go-Left versus Go-Right trials.

To evaluate the amount of task information in SC versus PFC populations, we used a cross-validated linear decoding approach (STAR Methods; Pagan et al., 2013). Although both populations contained above-chance task information throughout the trial duration, decoding for whether a trial was Pro or Anti was significantly more accurate in the SC than in the PFC for equally-sized populations (p<0.01, Figure 2C; n=193), even after controlling for firing rate differences between the two areas (Figure S3A). The stronger task information in the SC could only be matched when the number of neurons in a simulated PFC population was 5 times larger than the SC population (Figure S3B).

In addition, left versus right choice information appeared significantly earlier in SC than in PFC (latency difference = 191 ± 23 ms; p<0.01, Figure 2D). This choice information latency difference is not a result of firing rate differences between the two populations (Figure. S3A), and cannot be reduced by increasing the number of PFC neurons in a pseudo-population (Figure. S3C). These results argue against a model in which the decision is first computed in PFC and then relayed to SC. Instead, they suggest that the SC may play a role in forming the spatial decision.

### A subset of SC task-encoding neurons are linked to decisions

A closer examination of SC neurons revealed subpopulations that encode distinct types of information (Figure 3). For each SC neuron, we computed the temporal profile of significant task selectivity (Pro vs Anti), and ranked neurons by the time of their peak selectivity (Figure 3A). One group of SC neurons (which we labelled “cue neurons”, cyan) differentiated between Pro and Anti trials most strongly during the auditory cue, whereas another subpopulation represented task selectivity most strongly when the auditory cue was no longer present (“delay/choice neurons”, yellow). The representation of task context by cue neurons was progressively weakened after the end of the cue, and it did not differentiate between correct and error trials (Figure 3B, top), consistent with a purely sensory signal with little direct relationship to behavior.

**Figure 3.**
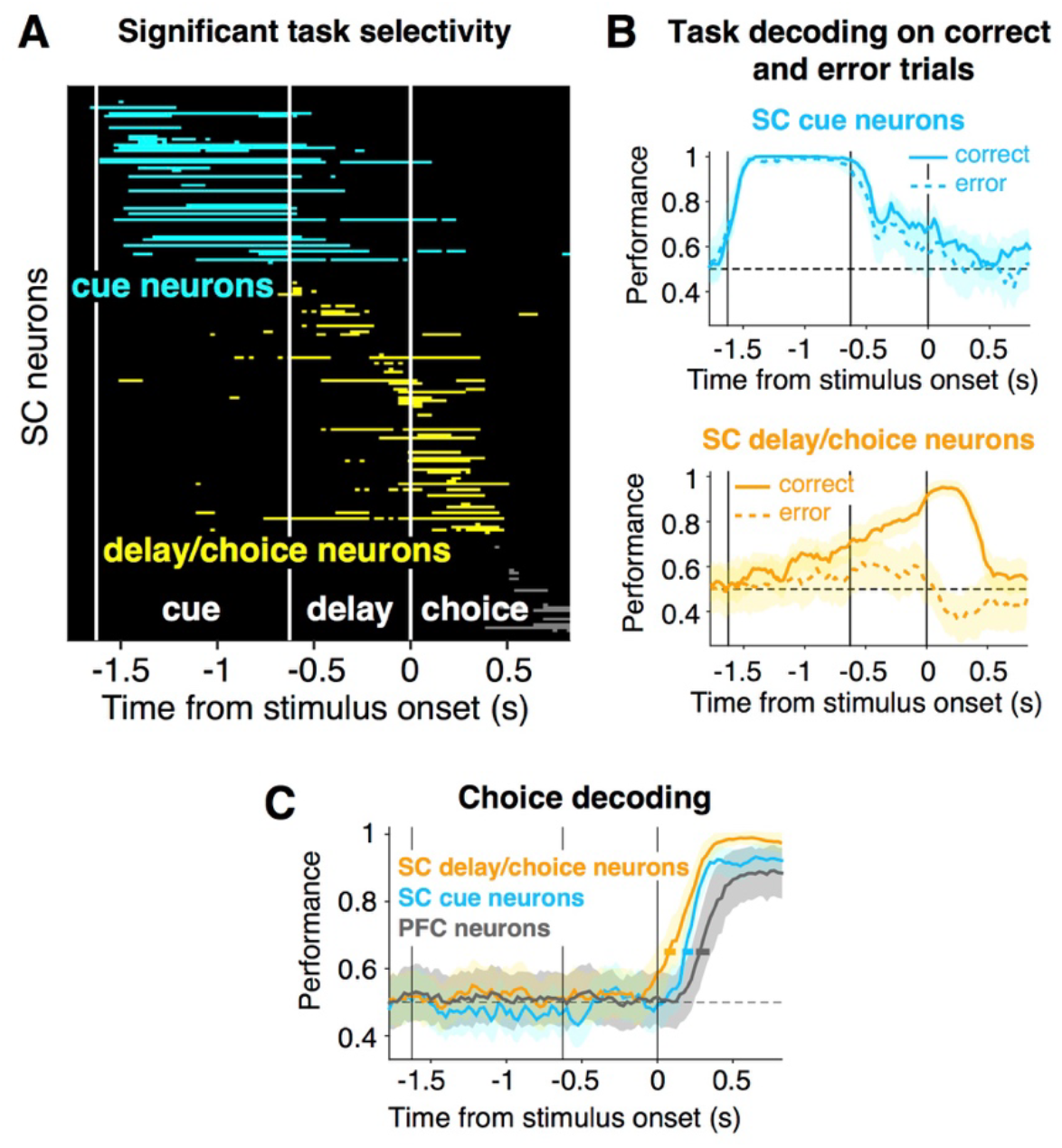
Distinct roles of SC subpopulations. (A) Timing of significant Pro/Anti selectivity (d’) for all SC neurons, sorted by peak d’. Significance threshold was determined by shuffled data. We separated SC neurons into two groups based on the timing of their Pro/Anti selectivity. “Cue neurons” (cyan, n=29) differentiated between Pro and Anti trials most strongly during the auditory cue; “delay/choice neurons” represented task selectivity most strongly when the auditory cue was no longer present (yellow, n=45). (B) Mean ± s.d. performance of task decoding on correct versus error trials. Linear classifiers trained on correct trials were tested for separate correct trials (solid) or error trials (dashed). The representation of task context by delay/choice neurons was disrupted on error trials whereas such information in the cue neurons did not differentiate between correct and error trials. (C) Choice decoding performance of SC subpopulations and PFC neurons (n=29 to match number of cue neurons, see STAR Methods). Choice information emerged first in SC delay/choice neurons. Shaded areas (vertical error bars) indicate s.d. of decoding accuracy for each population across time. Horizontal error bars represent s.d. of the timing of reaching 0.65 decoding accuracy for each population.

In contrast, three lines of evidence suggest that the delay/choice neurons play a key role in behavior. First, their task information slowly ramped up throughout the cue presentation and the delay to peak at the time when rats were required to make a motor choice (Figure 3B, bottom, solid line). Second, this representation was significantly disrupted on error trials during the delay and choice periods (Figure 3B, bottom, dashed line; p<0.01), indicating a strong correlation with behavior. Third, these neurons contained a very early representation of the correct choice, significantly faster than the SC cue neurons or the PFC neurons (Figure 3C, delay-choice neurons = 84 ± 19 ms, cue neurons = 196 ± 38 ms, PFC neurons = 290 ± 34 ms, p<0.01). Thus, the SC delay/choice neurons contain a performance-dependent task context signal (Pro vs Anti) that may be important for flexible routing of the upcoming target stimulus information (Left vs Right side light), so as to produce the context-appropriate orienting choice.

We therefore focused on the SC delay/choice neurons to examine how task and choice signals were multiplexed (Figure 4). In our behavior, the sensorimotor transformation occurs immediately after the visual target onset, when animals apply the non-spatial task context (Pro or Anti) to guide spatial orienting responses (ipsi- or contralateral to the recorded side). In contrast to the heterogeneity initially observed in the entire SC population (Figures 1C-E), focusing on the subset of delay/choice neurons during this critical time window revealed a systematic relationship between each neuron’s task selectivity and choice selectivity (Figures 4, S4B and S4C). Most neurons that fired more on trials with contralateral orienting responses also fired more during Pro trials (Figure 4A). Conversely, most Ipsi-preferring neurons were also Anti-preferring (Figure 4B). This suggests the existence of two broad groups of neurons during the choice period. We refer to these two groups of neurons as Pro/Contra-preferring neurons and Anti/Ipsi-preferring neurons. Note that this relationship between task and choice selectivity did not simply reflect a preference for the contralateral light stimulus, but revealed a complex interaction between task context information and target location (see example in Figure 1D, bottom).

**Figure 4.**
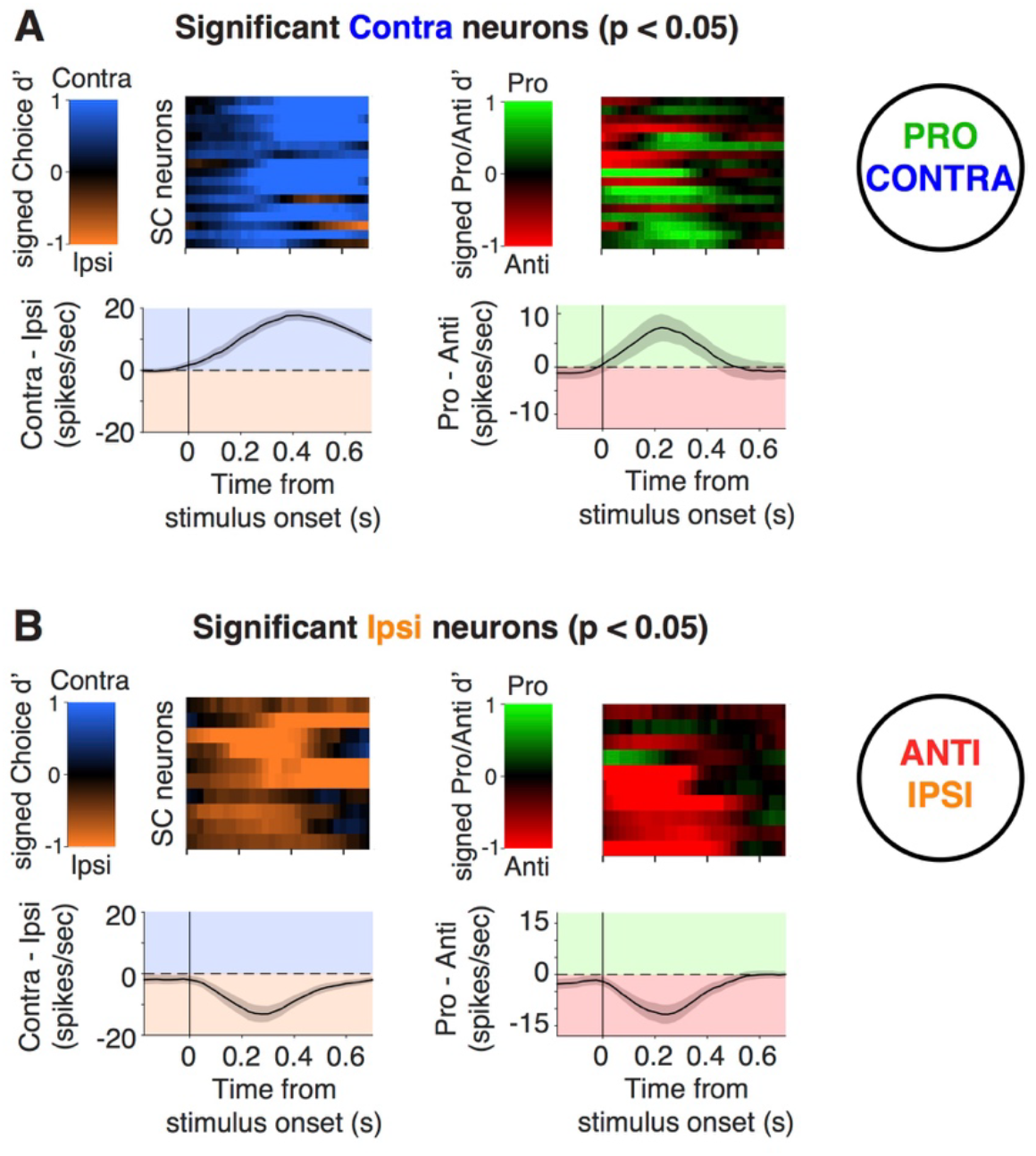
A relationship between task and choice encoding around stimulus onset, suggesting two groups of neurons. (A) Neurons selected as having a significantly greater firing rate on trials when the animal oriented contralaterally to the recorded neuron (n = 17). Left top shows one neuron per row, with the color indicating the strength of Contra/Ipsi selectivity (d’), as a function of time relative to the visual stimulus onset. Left bottom shows mean ± s.e.m. firing rate difference (Contra - Ipsi) averaged over these neurons. Right panels show the same neurons as in the left panels, but now analyzed for Pro/Anti selectivity and firing rate difference. Contra-preferring neurons tend to be Pro-preferring. (B) As in panel (A), but showing significantly Ipsi-preferring neurons (n = 10). Ipsi-preferring neurons tend to be Anti-preferring.

### SC activity is necessary during the non-spatial task-encoding delay period

Since different behavioral epochs of the task require distinct computations, we selectively probed the requirement of SC activity during separate epochs (Hanks et al., 2015; Kopec et al., 2015; Li et al., 2016) using bilateral optogenetic inactivation of SC neurons, mediated by virally-expressed eNpHR3.0, a light-activated chloride pump (Figures 5A, 5B and S2C, STAR Methods).

**Figure 5.**
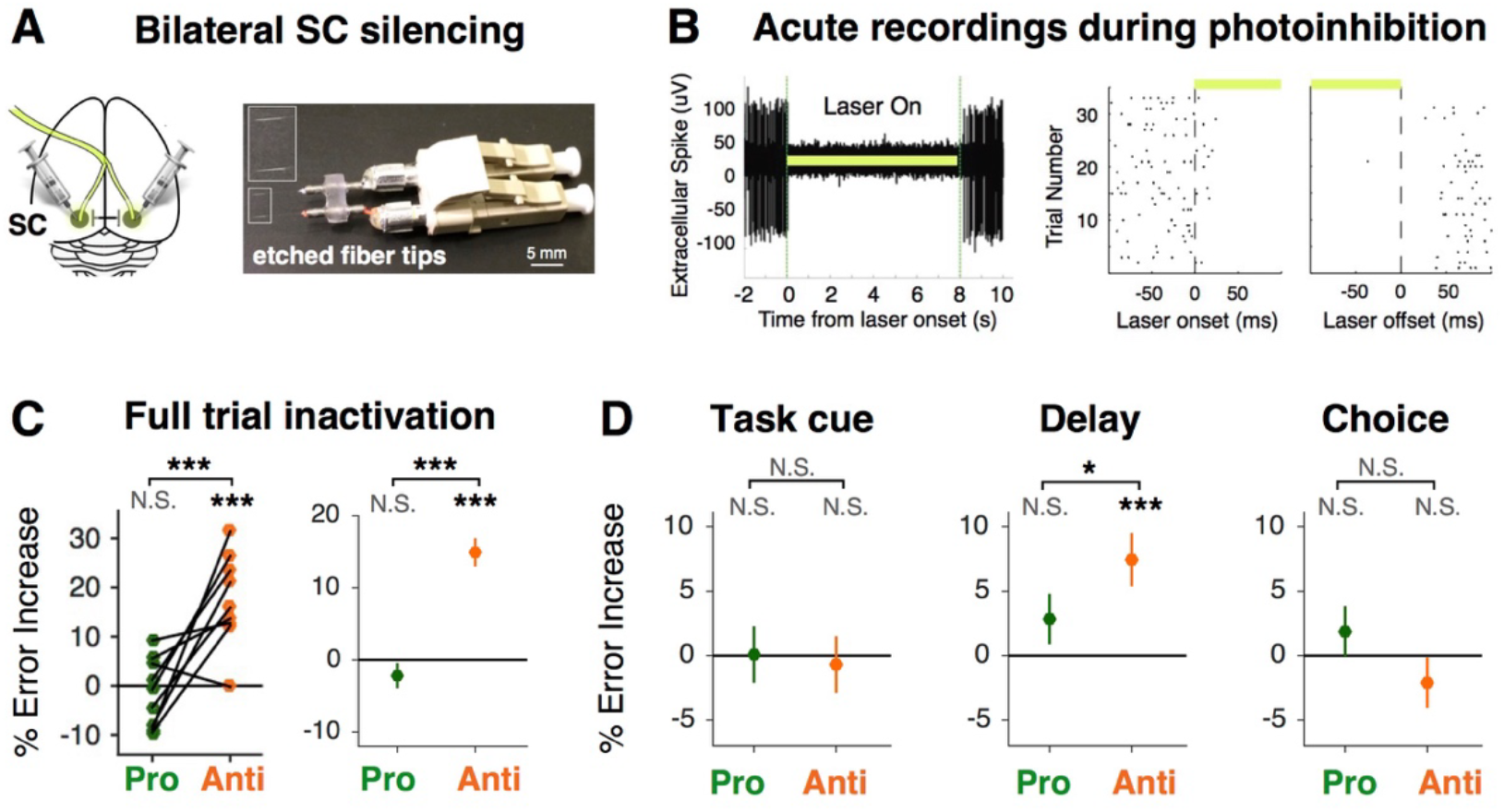
SC delay activity is required for the Anti task. (A) Experimental design for bilateral SC optogenetic inactivation. Left, a schematic that indicates virus infection and laser stimulation in the SC on both hemispheres. Right, an example of the optical fiber implant. The taper of each fiber is chemically sharpened to be approximately 2 mm long for stronger and more unified light delivery. The distance between the two fibers are constructed to be exactly 3.6 mm to target bilateral SC. (B) Physiological confirmation of optogenetic inactivation effect in an anesthetized animal. Left: acute extracellular recording of spontaneous activity in the SC expressing eNpHR3.0. Laser illumination period (8 s) is marked by the light green bar. Right: spike activity aligned to laser onset and laser offset over multiple trials. Note that the onset and offset of the inhibitory effect are on the scale of tens of milliseconds. (C) Effect of full-trial inactivation of bilateral SC. Mean Pro (green) and Anti (orange) error rate increase due to SC inactivation for all individual rats (n=9, left) and across all trials (Pro=662 trials, Anti=615 trials, right). Left, each data point represents the mean effect across sessions for a single rat. Right, means and s.e.m. across trials (concatenated across all 60 sessions). (D) Effect of sub-trial inactivations of bilateral SC on Pro and Anti error rate (mean and s.e.m. across trials from 102 sessions). Statistical comparison between Pro and Anti effects were computed using a permutation test, shuffled 5000 times. N.S. p>0.05; *p < 0.05; ***p < 0.001. Note that all types of inactivations were randomly interleaved for each session.

Optogenetic inactivation that covered the entire trial period (3 s) of a randomly selected 25% of trials resulted in a selective Anti impairment on those trials (Figures 5C and S5A; permutation test p<10-3 across animals or across all trials). This replicates previous pharmacological inactivation results where SC activity was suppressed during the entire session (Duan et al., 2015). Turning to temporally-specific inactivations, we found that bilateral SC inactivation during the task cue period did not result in any behavioral deficit (Figure 5D, left), consistent with a sensory role for cue neurons that are not required for correct performance (Figure 3B). In contrast, bilateral SC inactivation during the delay epoch significantly increased error rates on Anti trials (Figure 5D, middle; bootstrapped p<0.001). This suggests that an intact representation of task context, which is neither a spatial nor a motor variable, is required in the SC during the delay period in order for animals to perform the behavior. This finding is consistent with our electrophysiology data that linked delay neurons to flexible sensorimotor decisions (Figure 3B). Whether the task context representation is generated in the SC itself, or inherited from elsewhere, remains an open question. Finally, given the strong and early choice signal in the SC, we were surprised to find that bilateral choice period inactivation did not have any effect on choice accuracy (Figure 5D, right; p>0.05), although we did observe that correct Anti responses were slightly slowed down after choice period SC inactivation (RT increase = 22.5 ± 15.3 ms, p<0.05; Figure S5B).

### Collicular models of executive control consistent with perturbation data

Our electrophysiological results suggested grouping SC neurons as Pro/Contra or Anti/Ipsi. Could neural circuitry within the SC itself, between such Pro/Contra and Anti/Ipsi neuron groups, lead to the pattern of results seen in our optogenetic experiments (Figure 5D)? Or would choice formation circuitry external to the SC be necessary to explain the lack of a behavioral impairment following inactivation during the choice period? To address these questions, we developed a model framework in which the SC was represented by four pools of neurons, a Pro/Contra pool and an Anti/Ipsi pool, on each side of the brain (Figure 6A). Since unilateral SC stimulation drives contralateral orienting motions (Dean et al., 1988; Guitton et al., 1980; Robinson, 1972), we took the Pro/Contra neurons as driving the motor output, with the final choice determined by which of the two Pro/Contra units had greater activity (Goffart et al., 2012). The model had free parameters describing the sign and strength of connections between the units (Figure 6A), magnitude of additive noise, degree of silencing induced by optogenetic inactivation, and others (STAR Methods), for a total of 16 free parameters. Connections between the two sides of the SC can be both excitatory and inhibitory (May, 2006; Wolf et al., 2015), so we made no assumptions as to connection signs.

**Figure 6.**
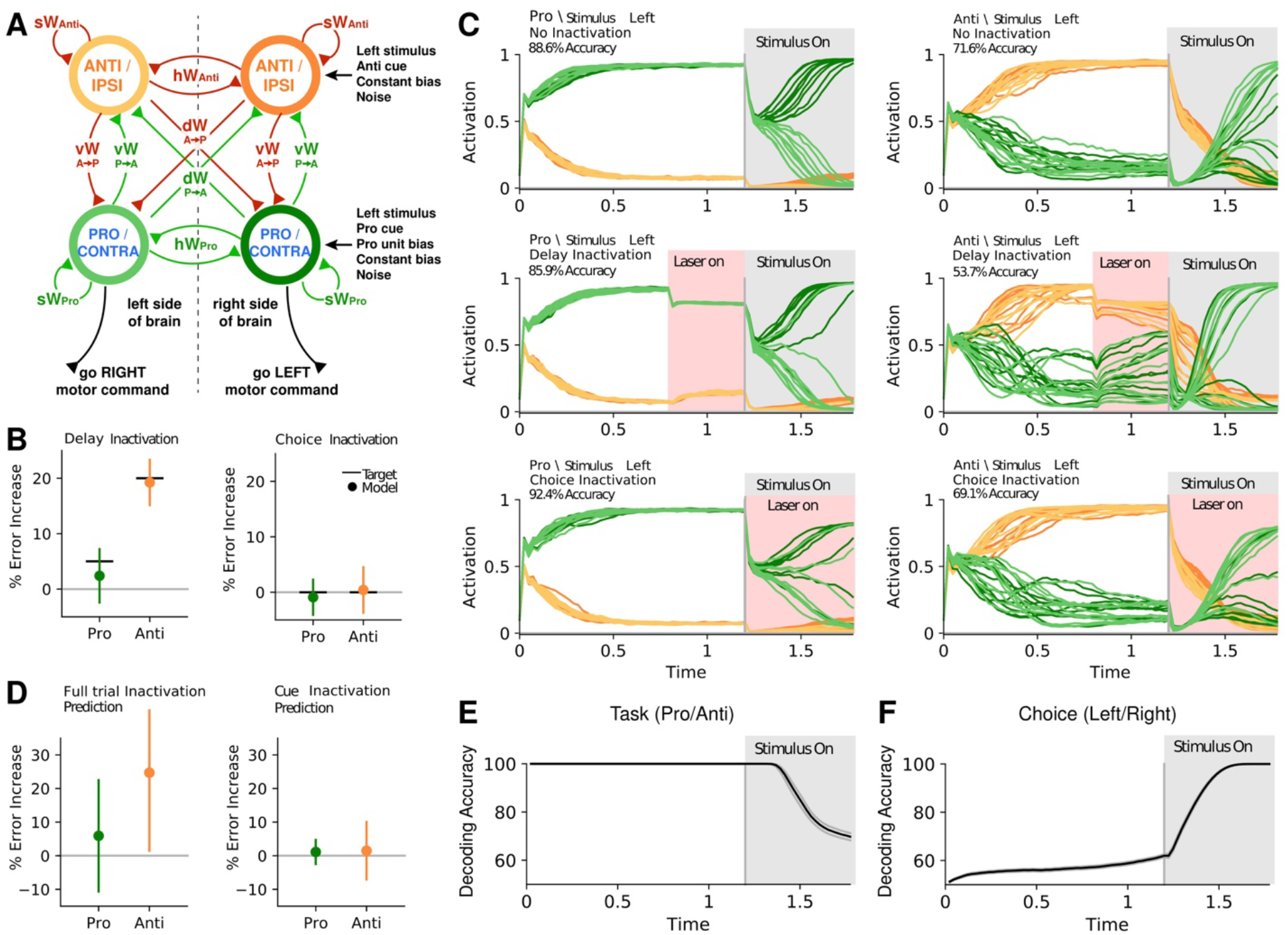
Dynamical SC model for flexible sensorimotor switching replicates optogenetic inactivation patterns. (A) Schematic of SC model, showing four units, each representing a population of SC delay/choice neurons. Connection strengths are independent in each direction, as well as for Pro and Anti self-connections, for a total of eight connection parameters. Each unit also receives external inputs. (B) Model fit to control, delay and choice period inactivation results. Target and model performance that qualitatively replicate the Pro (green) and Anti (orange) error rate changes after delay (left) and choice period (right) inactivations (mean ± 95% confidence intervals across all model solutions). (C) For one example solution, activity of the four SC units (colors as in A) on 10 example Pro (left) and Anti (right) trials in three different conditions: no inactivation (top row), delay period inactivation (middle row), and choice period inactivation (bottom row). Each trial had either 1 s or 1.2 s rule and delay period, follow by a 0.45 s or 0.6 s choice period. Light stimulus is presented on the left hemifield. Average accuracy on a test set of 10,000 trials is reported. Leftwards orienting responses correspond to trials where the dark green unit ends with higher activity than the light green unit, and rightwards orienting choices correspond to the opposite ordering. We do not assume that optogenetic inactivation in the experiments is necessarily complete; consequently, inactivations in the model (middle, bottom rows) reduce but do not completely silence activity. (D) Model prediction for full-trial inactivation (left) and task cue period inactivation (right), similar to (B). (E) Mean ± s.e.m. Pro/Anti task decoding (STAR Methods) across all model solutions as a function of time, correct and incorrect trials are included. (F) Mean ± s.e.m. Left/Right choice decoding across all model solutions as a function of time, correct and incorrect trials are included.

We optimized model parameters to qualitatively match the optogenetic inactivation results: choice period inactivation had no effect on choice accuracy (Figure 5D, right); delay period inactivation impaired accuracy on Anti trials, but not Pro trials (Figure 5D, middle); and control trials had higher accuracy for Pro compared to Anti (Figure S1C; Duan et al., 2015). We performed the optimization starting from many random parameter values. We refer to the 373 optimizations with low final error as solutions (STAR Methods). When each solution was then tested with a different noise realization, the models replicated the inactivation patterns we observed in experimental data (Figures 6B and 6C). Model solutions that fit the delay and choice period inactivation data can also predict the full-trial and cue-period inactivation data, with variability across solutions (Figure 6D). To investigate whether the model dynamics encode similar information as those found in recorded SC neurons, we decoded Pro/Anti task and left/right choice for each model solution (STAR Methods). The average decoding patterns parallel those found in our electrophysiology data (Figure 2): rule information during the task cue and delay period (Figure 6E), followed by a choice signal after target presentation (Figure 6F). We conclude that collicular circuitry involving Pro/Contra and Anti/Ipsi neurons is sufficient to reproduce baseline Pro/Anti accuracies, a lack of a choice effect during choice period silencing, and selective impairment of Anti trials during delay period silencing.

We next sought to uncover the mechanisms that each solution uses to solve the task. We used singular value decomposition (SVD) to find a low dimensional representation of all solutions in terms of their dynamics (STAR Methods). To our surprise, the dynamics across solutions were highly heterogeneous (Figure 7A). Example trials from several representative solutions reveal different rule encoding strategies (Figure 7B). The variability in dynamics is also reflected in variability in parameter values (Figure S6). Principal Components Analysis on the parameter values or dynamics finds that more than 10 principal components are required to explain 90% of the variance in parameter values or dynamics across the 373 solutions (Figure S7A), even though fewer than 3 dimensions were needed to explain variance for dynamics within each solution (Figure S7B). These findings demonstrate that this simple dynamical circuit can perform the task using a large variety of configurations.

**Figure 7.**
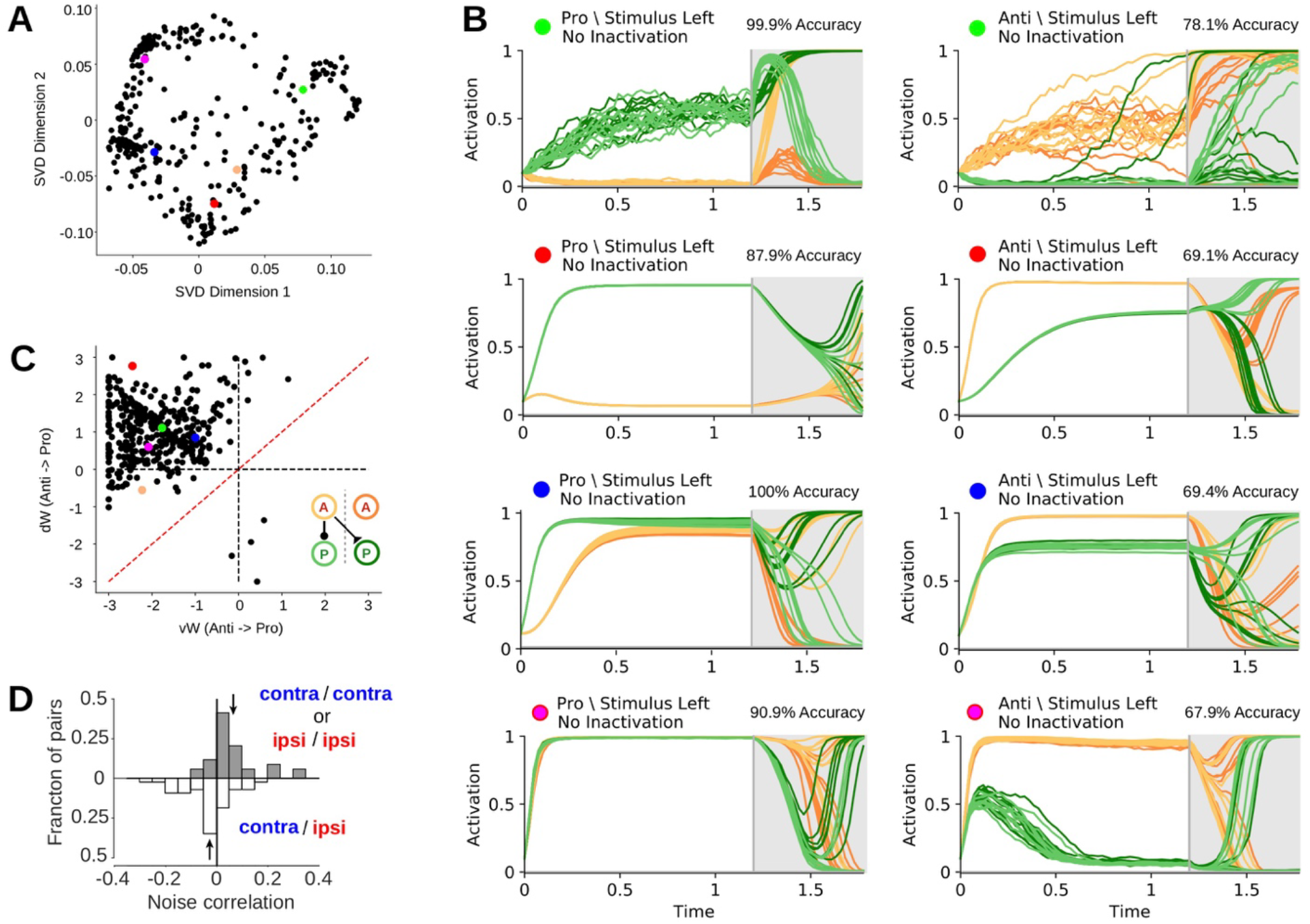
Common circuit mechanisms across heterogeneous model dynamics. (A) Projection of model solutions onto the top two dimensions that maximally explains variance in model dynamics across solutions. Individual dots are unique model solutions. Colored dots refer to the example solutions shown in (B). Orange dot is the example in Figure 6. (B) Model unit activity on 10 Pro and Anti trials from 4 example solutions, similar to Figure 6C (control only). (C) Scatter plot of diagonal weights (from Anti units to Pro units on the opposite hemisphere) against vertical weights (from Anti units to Pro units on the same hemisphere), for all model solutions. Equality line added for reference. Colored dots refer to example solutions shown in (B). Insert cartoon summarizes mean connectivity patterns with inhibition from the Anti unit to the ipsilateral Pro unit, and excitation to the contralateral Pro unit. (D) Trial-by-trial noise correlation between pairs of simultaneously recorded neurons on one side of the SC, calculated for within-group pairs of neurons (Contra/Contra or Ipsi/Ipsi, upper histogram) and between-group pairs (Contra/Ipsi). Noise correlation distribution for within-group pairs was significantly shifted above 0 (mean = 0.082 ± 0.02; p<0.01), whereas the between-group distribution was significantly shifted below 0 (mean = −0.052 ± 0.02; p<0.05), consistent with negative vertical weights as predicted by the model. Arrows indicate the mean values.

Despite the highly varied dynamics and parameter values in different solutions, we identified two features that were consistent across model solutions. First, almost all solutions had inhibitory connections from the Anti unit to the Pro unit on the same hemisphere (vertical weight from Anti to Pro unit, 365/373 negative, 97.9%, Figure 7C). This suggests that suppression of the “default” Pro pathway by Anti task representation locally in the SC may be essential for avoiding reflexive Pro responses towards the target stimulus. Consistent with this modeling result, noise correlations between pairs of simultaneously recorded SC neurons encoding opposite task contexts were significantly negative (Figure 7D). Note that the connections from the Pro unit back to the ipsilateral Anti unit were not necessarily inhibitory (vertical weights from Pro to Anti unit, only 252/373 negative, 67.6%, Figure S6), indicating an asymmetry between Pro and Anti task representation in the circuit architecture that parallels behavior. Second, most solutions had excitatory connections from the Anti unit to the Pro unit on the opposite hemisphere (diagonal weights from Anti to Pro unit, 339/373 positive, 90.9%, Figure 7C), constituting the pathway that executes the task-appropriate orienting response *away* from the target stimulus during Anti trials. For the small fraction of solutions with positive vertical weights from the Anti to the ipsilateral Pro unit (dots to the right hand side of vertical dashed line in Figure 7C) or negative diagonal weights (dots below horizontal dashed line in 7C), most of them (34/38, 89.47%) had a more positive projection from the Anti unit to the contralateral Pro than to the ipsilateral Pro unit, consistent with a functionally competitive inhibition mechanism between the two projections. Across all solutions, 369/373, 98.9% are above the dashed red line (Figure 7C).

Thus, by optimizing model solutions to match behavioral and optogenetic results, we found anatomical and functional features of a circuit, modeled as lying within the SC itself, that correspond to key processes of executive control: 1) inhibiting task-inappropriate behavior and 2) executing task-appropriate actions. These common features reflect fundamental principles that could constrain circuit behavior, and support an executive control function for the superior colliculus.

## DISCUSSION

Past studies of executive control have focused on the prefrontal cortex (PFC) as the “command center” that either biases downstream areas to achieve internally-generated goals (Miller and Cohen, 2001) or directly inhibits other regions to suppress context-inappropriate actions (Munoz and Everling, 2004; Pierrot-Deseilligny et al., 1991a). In contrast, the midbrain superior colliculus (SC), traditionally considered a spatiomotor structure, has been hypothesized to mediate fast, stimulus-driven responses (Everling et al., 1998), rather than flexible control of behavior. Our analysis of prefrontal and midbrain activity during flexible sensorimotor routing provides three lines of evidence for an extended executive control network that includes the SC. First, SC neurons contain stronger non-spatial task context information and earlier decision information than PFC neurons (Figures 1, 2 and 3). Second, an intact task context representation within the SC itself is causally required for behavior (Figure 5). Finally, our computational models demonstrate that fast sensorimotor routing can be achieved through control of nonlinear dynamics, within collicular circuits themselves, in a manner consistent with our behavioral, electrophysiological, and optogenetic findings (Figures 4, 6, and 7).

Decision signals for the animal’s spatial orienting response were found to emerge early in the SC, less than a hundred milliseconds after the side light was turned on and the spatial choice could begin to be computed. The latency for these signals in the SC was far shorter than in the PFC, by a difference of ~200 ms (Figure 3C), suggesting that the motor choice output of flexible sensorimotor transformations is not first computed in the PFC, but is instead first computed in a circuit involving the SC. We note that the SC, which is part of the midbrain, is a highly conserved structure of the vertebrate brain, whereas the prefrontal cortex has undergone dramatic changes during evolution (Preuss, 1995; Preuss and Kaas, 1999; Seamans et al., 2008; Uylings et al., 2003); this underscores an important question left open, namely whether or not the relative roles of the midbrain and prefrontal regions in executive control are conserved across rodents and primates. We have structured our behavior to closely parallel analogous primate paradigms with the goal of facilitating such cross-species analysis. Single-neuron recordings in monkey SC (Chan et al., 2017; Everling et al., 1998, 1999) and dorsolateral PFC (dlPFC; Everling and DeSouza, 2005; Johnston and Everling, 2006a, 2006b; Johnston et al., 2009) have revealed heterogeneous task-relevant signals comparable to those observed here (Figure 1), but a direct comparison of monkey dlPFC and SC population decoding accuracy and latency, so as to test the relative strength and timing of their decision signals, or temporally-specific silencing of these structures, have yet to be conducted in monkeys.

The causal contribution of SC activity to spatial decision-making, spatial target selection, and spatial attention has mainly been investigated using unilateral perturbations (Felsen and Mainen, 2008; Kopec et al., 2015; McPeek and Keller, 2004; Song et al., 2011; Stubblefield et al., 2013; Zénon and Krauzlis, 2012). These experiments, all within the spatial domain, have established the SC’s importance in contralateral control of orienting responses. Consistent with those findings, unilateral pharmacological inactivation of the SC during our ProAnti task also resulted in a contralateral impairment (Duan et al., 2015). Here, we used bilateral perturbations. These revealed a selective requirement of SC activity for the cognitively demanding Anti task during the delay period (Figure 5D), dissociated from motor planning or spatial decision processes. These results indicate that the active representation of relevant, non-spatial task goals, locally within the SC, plays a key role in flexible sensorimotor routing.

To gain mechanistic insights regarding a collicular circuit for executive control, we turned to dynamical models that were inspired by our electrophysiology data (Figure 4) and constrained by the optogenetic results (Figure 5). It has been observed that as proposed circuit models in biology grow in complexity, even high-throughput experiments may provide data that constrain only a fraction of the characteristics that fully define the circuit, and human intuition may prove insufficient to find the full space of possible solutions (Fisher et al., 2013; Goldman et al., 2001; Gutierrez et al., 2013; Prinz et al., 2004; Transtrum et al., 2015). Here, we used computational methods to find a wide variety of SC models that were compatible with our behavioral and optogenetic inactivation data (Figure 6). These results demonstrate the sufficiency of collicular circuitry involving physiologically identified neuron subtypes to implement flexible sensorimotor routing.

By examining common features across distributions of successful yet highly varied solutions, we identified potential intra-collicular circuit mechanisms for the two fundamental processes of executive control: suppression of automatic responses and vector inversion (Figure 7; Munoz and Everling, 2004). First, we found that the inhibition of the “default” Pro pathway by Anti representation on the same side of the SC was a key component for almost all model solutions to achieve good performance (Figure 7C). Such inhibitory control mechanism has previously been proposed as an important aspect of executive control to suppress context-inappropriate actions (Lo and Wang, 2016; Munoz and Everling, 2004; Pierrot-Deseilligny et al., 2003), but via prefrontal cortical inhibition of SC activity (Everling et al., 1998; Pierrot-Deseilligny et al., 1991a). However, a recent monkey pro/antisaccade study showed that the primary influence of dorsolateral PFC neurons on the SC was not inhibitory but excitatory (Johnston et al., 2014). Our experimental and modeling results argue that response suppression may be carried out, at least in part, by local SC neurons that represent task context themselves.

Second, we found that model Anti units activated Pro units on the opposite hemisphere, to implement the Anti orienting responses away from the target stimulus (Figure 7C), hypothesizing a novel modulatory function for the excitatory SC neurons that project to the contralateral output neurons (May, 2006; Wolf et al., 2015). Future experiments that combine projection-based electrophysiology (Economo et al., 2018; Li et al., 2015) with immunohistochemistry can test these predictions to further constrain the circuit logic for context-based action selection. The competitive inhibition between the left and right Pro/Contra units via Anti unit activation also provides an alternative mechanism to determine decision output. This is often postulated as being driven by direct mutual inhibition between the two output units themselves (Machens et al., 2005; Wong and Wang, 2006), but we did not find in our models that direct mutually inhibitory connections between the two output units were a necessary feature for most solutions (horizontal weights between Pro units can be either excitatory or inhibitory, Figure S6). It should be noted that our task only involves two choice options. Whether the SC is important for vector inversion in a more general sense, involving continuous variables, remains to be tested with a continuous choice set.

Together, our experimental and modeling work suggest that cognitive functions that are normally associated with PFC circuit (Mante et al., 2013; Miller and Cohen, 2001; Rigotti et al., 2013) could be equally attributed to the midbrain SC (Crapse et al., 2018; Herman et al., 2018; Horwitz et al., 2004; Zénon and Krauzlis, 2012). The SC motor layers receive projections from virtually the entire brain (Edwards et al., 1979; May, 2006; Sparks and Hartwich-Young, 1989), forming multiple cortico-subcortical loops that are well-situated for cognitive functions such as rapid sensorimotor routing and flexible control of instinctual behavior (Evans et al., 2018, 2019; Yilmaz and Meister, 2013). Whether PFC and SC represent nodes of task context representation in parallel circuits (Zénon and Krauzlis, 2012) or part of the same recurrently-connected network (Kopec et al., 2015) remains to be further investigated. Our results call for a broadening of focus in basic and clinical studies of executive functions to include interconnected cortical and subcortical areas (Basso and May, 2017; Everling and Johnston, 2013; Krauzlis et al., 2013; Schmitt et al., 2017; Wimmer et al., 2015).

## STAR METHODS

### Subjects

Nineteen adult male Long-Evans rats (Taconic) were used for the experiments presented in this study. Of these, 7 rats were used for electrophysiology recordings, and 12 rats were implanted with optical fibers for the optogenetic inactivation and YFP control experiments. Animal use procedures were approved by the Princeton University Institutional Animal Care and Use Committee and carried out in accordance with NIH standards.

### Behavior

Rats were trained on the ProAnti task-switching behavior (Duan et al., 2015). Each trial began with an LED turning on in the center port, instructing the rats to nose poke there to initiate a trial. They were required to keep their noses in the center port until the center LED offset (nose fixation). Broken fixation trials were ignored in all analyses. During the first 1 s of nose fixation, a Pro or Anti sound was played (clearly distinguishable FM modulated sounds) to indicate the current task, followed by a 500-ms silent delay when rats had to remember the current task while maintaining nose fixation. The center LED was then turned off, allowing the animal to withdraw from the center port. The withdrawal would trigger either a left or right LED to turn on as the target stimulus, which remained on until rats poked into one of the side ports. Response time (RT) is defined as the time from target onset until side poke. On a Pro trial, rats were rewarded for orienting towards the side LED; on an Anti trial, rats were rewarded for orienting away from the side LED and into the port without light. A correct choice was rewarded by 24 μl of water; and an incorrect choice resulted in a loud sound, no reward, and a short time-out. To ensure that all sub-trial optogenetic inactivation conditions have the same duration for laser stimulation (750 ms), all rats implanted with optical fibers were trained on a modified version of the behavior where the task cue period and the delay period both lasted 750 ms instead of the 1-s cue period and the 500-ms delay period as in the original design.

In all recording and inactivation sessions, rats performed alternating blocks of Pro and Anti trials, where block switches occurred within single sessions, after a minimum of 15 trials per block, and when a local estimate of performance (over the last ten trials in this block) reached a threshold of 70% correct. Detailed training procedures and codes can be found in a previous report (Duan et al., 2015).

### Recordings

Rats were implanted with custom-made movable microdrives and recordings were made with platinum-iridium tetrodes (Erlich et al., 2011). To target the prelimbic (PL) area of PFC (+3.2 anteroposterior [AP] mm, ±0.75 mediolateral [ML] mm from bregma), tetrodes were initially positioned at ~1.5 mm below brain surface and were advanced daily during recording sessions to sample different neurons. To target the intermediate and deep layers of the SC (−6.8 AP mm, ±1.8 ML mm), tetrodes were initially positioned at ~3 mm below brain surface and advanced daily. Electrode placements were confirmed with histology. Four rats had both PL and SC implants (same hemisphere), 2 rats had a PL implant only, and 1 rat had an SC implant only. The choice of recording area and hemisphere side was assigned randomly for each rat.

### Analysis of neural data

Spike sorting was done manually using SpikeSort3D (Neuralynx), and only isolated single units were included in the following analyses. In order to perform analyses on the neural population, we only analyzed neurons recorded for a sufficient number of trials. More specifically, we only analyzed neurons for which we had collected responses during at least 25 correct trials for each of the four possible task conditions (Pro-Right, Pro-Left, Anti-Right, Anti-Left). This resulted in the analysis of 193 neurons (out of 215) in SC, and 291 neurons (out of 331) in PFC. Unless otherwise noted, all analyses were performed on correct trials. The response of each neuron was quantified by counting the number of spikes in 250ms-wide bins. In all analyses, the response was aligned to the time when the target stimulus appeared (i.e. the time of withdrawal from the center port). The temporal gap between the fixation offset and target stimulus onset was controlled by animals and thus variable on each trial. On average, rats withdrew from the center port 127 ms after fixation offset. Therefore, in all figures, we indicate the start of the delay period (end of task cue presentation) 0.627 s before target stimulus onset (500 ms delay + 127 ms), and the start of task cue presentation at 1.627 s before target onset (1 s of task cue presentation before the delay). Unless otherwise noted, in all figures the time scale is causal, i.e. the value at time 0 refers to the neural activity in a time bin between −250 ms and 0 ms.

## Quantification of single neuron selectivity

The amount of information encoded by a single neuron about a task variable was measured at each time point using d’, defined as the difference in the number of spikes fired in response to two generic task conditions (here named as A and B), normalized by the square root of the pooled variance: 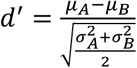, where *μ*_*A*_ indicates the mean spike count in response to condition A, *μ*_*B*_ indicates the mean spike count in response to condition B, *σ*^2^_*A*_ indicates the variance across trials of the spike count in response to condition A, and *σ*^2^_*B*_ indicates the variance across trials of the spike count in response to condition B.

Information about the task rule (Pro/Anti d’) was computed by comparing the responses during Pro trials and the responses during Anti trials (with positive d’ indicating Pro-preference). Information about the rat’s choice (Choice d’) was computed by comparing the responses during trials that resulted in an orienting movement contralateral to the recorded neuron, and trials that resulted in an ipsilateral orienting movement (with positive d’ indicating Contra-preference; Figures 1C-H).

The threshold above which a d’ value was considered significantly different than 0 was computed based on the pairwise t-test between the two conditions, using a p-value of 0.05. In Figure 3A, d’ significance at each time point was computed using a shuffling procedure to correct for multiple comparisons, where the d’ at each time point was recomputed 100 times after randomly shuffling the labels of Pro and Anti trials, and the 95th percentile of the resulting overall distribution of shuffled d’s was used as the significance threshold.

Single neuron selectivity about the task rule was used to define two distinct classes of neurons (Fig. 3A). “Cue neurons” were defined as those with peak Pro/Anti d’ at a time while the task cue was still being presented. “Delay/Choice neurons” were defined as those with peak Pro/Anti d’ at times after the task cue was no longer present. Neurons whose Pro/Anti d’ was never significantly higher than 0 were excluded from both groups.

Within the class of “Delay/choice neurons”, we used single neuron selectivity about the choice in the first time bin after stimulus presentation (i.e. from 0 to 250 ms) to further subdivide these cells into two groups (Fig. 4). “Contra neurons” had a significantly higher response to contralateral stimulus, whereas “Ipsi neurons” had a significantly higher response to an ipsilateral stimulus.

### Population-level decoding analysis

To determine the amount of task-relevant information available in the SC and PFC neural populations at each time point, we performed a series of cross-validated linear classification analyses (Pagan et al., 2013). For each analysis, we considered the spike count responses of a population of N neurons to a task condition as a population “response vector” **x**, and we randomly assigned 60% of the recorded trials (30 trials) as the training set, and the remaining 40% of the trials (20 trials) as the test set. The training set was used to compute the linear hyperplane that would optimally separate the population response vectors corresponding to two different task conditions (e.g. Pro trials vs Anti trials). This linear readout can also be written as *f*(***x***) = ***w***^*T*^***x*** + *b* where **w** is the N-dimensional vector of weights applied to each of the neurons, and b is a scalar threshold. The classification of a test response vector **x** was then assigned depending on the sign of f(**x**), and the performance was computed as the fraction of correct classifications over 500 resampling iterations. Because some of the neurons were recorded in different sessions, trials were always shuffled on each iteration to destroy any artificial trial-by-trial correlations. The hyperplane and threshold were computed using a Support Vector Machine algorithm using the LIBSVM library (https://www.csie.ntu.edu.tw/~cjlin/libsvm).

When comparing the classification performances for neural populations with different numbers of neurons, we randomly resampled identical numbers of neurons without replacement on each iteration. Because the overall average firing rate was higher in SC than in PFC, we tested whether matching firing rates was sufficient to explain the classification result (Figure S3A), by removing single spikes at random from the SC dataset until the average firing rates were matched, and by performing again the classification analysis on the equalized SC population. To produce an estimate of the number of PFC neurons necessary to match performances in SC (Figure S3B), we adopted an analytical approach to estimate classification performances based on the distribution of d’ in a neural population (Pagan and Rust, 2014a). More specifically, we quantified the Normalized Euclidean Distance (NED) between two conditions in a neural population as the square root of the sum of the squared d’s across all neurons: 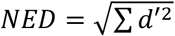. This quantity can be used to estimate linear classification performances, under the Gaussian assumption, as^48^: 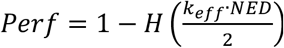, where H is the complementary error function, and *k*_*eff*_ is an efficiency factor that accounts for the inability of the classifier to extract all the available information (e.g. due to limited training data). Before applying this formula, d’s had to be corrected for their intrinsic positive bias (Pagan and Rust, 2014b). Because the NED grows with the square root of the total number of neurons in a population, we could then estimate the classification performance for a neural population of arbitrary size M as: 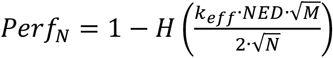, where N indicates the actual size of the population for which NED was computed. Using the same approach, we also tested how measurements of latency in the rise of classification performance (see below) depended on the total number of neurons in a neural population (Figure S3C).

When classification analyses were used to compare performances during correct and error trials (Figure 3B), we always trained the classifier using correct trials, and we tested the classifier using either correct or error trials. The number of trials used for testing was limited by the neuron with the fewest number of error trials per condition (9 trials).

To compute the latency of the rise in choice classification performance for different neural populations (Figure 3C), we evaluated the average time after the appearance of the target stimulus necessary for the population readout to reach a fixed threshold (correct performance >65%) (Pagan and Rust, 2014a). More specifically, on each iteration of the resampling procedure we computed the classification performances for each time point, we smoothed the resulting curve by averaging the value for 5 neighboring time points, and we noted the time point where the curve crossed the performance threshold. We computed the mean and the standard error of the latency as the mean and standard deviation of these values.

To compute the significance of differences in the magnitude (or latency) of population performances, we adopted a bootstrap approach based on our resampling procedure (Efron and Tibshirani, 1993). More specifically, we first evaluated the average performance (or latency) across all iterations for the two populations, and we then computed the p-value as the fraction of iterations in which, by chance, the value for the population with the lower average was above the value for the population with the higher average.

### Optical fiber construction, virus injection and fiber implantation

Chemically sharpened optical fibers (50/125 um LC-LC duplex fiber cable, http://www.fibercables.com) were prepared as previously described (Hanks et al., 2015). To ensure the distance between the two optical fibers was the distance between bilateral SC (3.6 mm), we inserted two metal cannulae into a plastic template and guided the optical fibers through the cannulae, which were 3.6 mm apart (Figure 5A).

Basic virus injection techniques were identical to those described previously (Hanks et al., 2015). At the targeted coordinates (SC, −6.8 AP mm, ±1.8 ML mm from bregma), two injections of 9.2 nl AAV virus (AAV5-CaMKIIα-eYFP-eNpHR3.0 for inactivations, 9 rats; AAV5-CaMKIIα-eYFP for controls, 3 rats) were made every 100 um in depth starting 3.5 mm below brain surface for 1.5 mm. Four additional injection tracts were completed, one 500 um anterior, posterior, medial, and lateral from the central tract. A total of 1.5 μl of virus was injected over the course of 30 minutes. Chemically sharpened bilateral SC fiber implant was lowered down the central injection track, with the tip of each fiber positioned at 4.4 mm below brain surface to target the center of SC’s intermediate and deep layers. Training was resumed 5 days post-surgery. Virus expression was allowed to develop for 8 weeks before behavioral testing began.

### Optogenetic inactivation and analysis

For each inactivation session, animals’ implants were connected to a 1m patch cable connected to a fiber rotary joint (Princetel) mounted above the behavioral chamber. A 200 mW 532 nm laser (OEM Laser Systems) was then connected to deliver constant light at 25 mW per site, with a < 5 mW difference between the left and right SC. Laser illumination occurred on 25% randomly chosen trials in each behavioral session. Different optogenetic conditions (3-s full-trial inactivation, 750-ms task cue, 750-ms delay, or 750-ms choice period inactivation) were randomly interleaved for all sessions to control for behavioral fluctuations across days.

Behavioral changes due to optogenetic inactivation were quantified as the performance difference between inactivation (laser) trials and control (no-laser) trials from the same sessions. These results are then compared to YFP control data. For each session, we calculated the baseline error rate or RT for Pro and Anti control trials and subtracted that mean value from the performance on individual inactivation trials. After obtaining the normalized changes in performance due to inactivation for individual sessions, we concatenated trials across all sessions and all rats, and computed the mean and s.e.m. across trials. Nonparametric bootstrap procedures or permutation tests were used to compute significance values (shuffled 5000 times). All rats were included in the full-trial inactivation analyses. For sub-trial inactivation analyses, we only included the rats (8/9) that had significant full-trial effects.

### Acute characterization of optogenetic effects

To measure the effects of optogenetic inactivation on neural activity, acute recordings of infected SC neurons were performed in anesthetized rats (Figure 5B). An etched fiber optic and sharp tungsten electrode (0.5 or 1.0 MΩ) were independently advanced to the center of the infected area. For each neuron tested, baseline neural activity was recorded for 2 s, followed by 8 s of laser stimulation at 25 mW, and another 2 s of post-stimulation recording, repeated for >10 times. We observed that the onset and offset of optogenetic inactivation of neural activity was within 50 ms of laser onset and offset (Figure 5B).

### Model setup

Our model consists of four dynamical units, each unit had an external(*V*_*i*_) and internal (*U*_*i*_) variable. The relationship between the internal and external variables is given by:

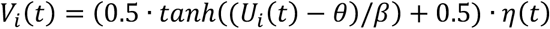

Here *η*(*t*) is the optogenetic inactivation fraction, which tells us the fraction of this unit’s output that is silenced by optogenetic inactivation in a time-dependent fashion (1 = no optogenetic inactivation). *β* = 0.5 controls the slope of the input-output relationship, and *θ* = 0.05 controls the midpoint of the input-output function. The internal variables had dynamical equations:

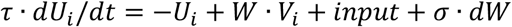

Where *W* is the network weight matrix, *input* is the external input into the network, *τ* = 0.09 *s* is a fixed time constant for each unit, and *σ* · *dW* is gaussian noise with amplitude given by the parameter *σ*.

*W* was parameterized by eight parameters that controlled the Pro self-weights *sW_P*, and Anti self-weights *sW_A*, the horizontal weights *hW_P* between the two Pro units and *hW_A* between the two Anti units, the vertical weights *vW_PA* from the Pro unit to the Anti unit on the same side and *vW_AP* from the Anti unit to the Pro unit on the same side, and the diagonal weights *dW_PA* from each Pro unit to the Anti unit on the opposite side and *dW_AP* from each Anti unit to the Pro unit on the opposite side. The external input into the network was given by:

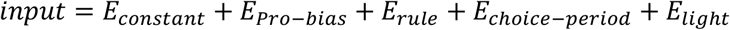

*E*_*constant*_ is constant excitation to all units. Parameter *E*_*Pro-bias*_ is constant excitation to both Pro units, but not to the Anti units. *E*_*rule*_ is the rule input, which is only active during the rule and delay periods, and not during the choice period. On Anti trials, the two Anti units get rule input *E*_*Anti-rule*_ and on Pro trials, the two Pro units get rule input *E*_*Pro-rule*_. Parameter *E*_*choice-period*_ is excitation to all units only during the target period when a light cue is presented and animals are free to choose. Parameter *E*_*choice-period*_ is excitation to both units on the side (L vs R) activated by the light cue, when the cue is active. Each trial was simulated numerically used the forward-euler method with time step dt=0.024s, which we found to balance accuracy and computational speed. To encourage robust solutions, we trained the network on four different trial lengths. The rule + delay period was either 1s or 1.2s, and the target period was either 0.45s, or 0.6s. Individual trials of the same trial type and duration are differentiated by the noise samples generated by the additive gaussian noise process.

### Model cost function

The cost function has two terms *C* = *C*_1_ + *C*_2_, *C*_1_ penalizes model performance that deviates from the target performance, *C*_2_ penalizes weak model choices where the output units are close together. Below we will describe *C*_1_ and *C*_2_ respectively.

#### *C*_1_ *Term*

To read out the models choice on a given trial, we could ask if *V*_*Pro-R*_ > *V*_*Pro-L*_. However, this creates a discontinuity in the cost function if a small change in a parameter causes the decision to flip. In order to use powerful optimization tools like automatic differentiation, we wanted the cost function to be fully differentiable. Therefore, each model choice was recast as the probability of a choice by passing the unit outputs through a tanh() function with a sensitivity given by a fixed parameter *θ*_1_. For a Pro trial, the probability of a correct choice was given by: HitP =0.5 * (1 + *tanh*((*V*_*Pro-R*_ − *V*_*Pro-L*_)/*θ*_1_)), and for an anti trial: HitA = 0.5 * (1 + *tanh*((*V*_*Pro-L*_ − *V*_*Pro-R*_)/*θ*_1_)). For each trial type, *i*, we defined a target hit percentage, and penalized the difference between the target hit percentage and the average hit percentage from the model across all trials. The overall cost from this first term was the sum across trial types. 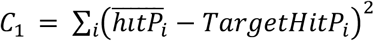.

#### *C*_2_ *Term*

The tanh() makes the cost function differentiable, but encourages the model to reach the target hit percentage on *every trial*, rather than making strong choices on each trial, some right and some wrong, that average to the target hit percentage. To prevent this degenerate solution, we introduced a second cost term that penalizes weak choices where the activation of the two Pro units are close. For a Pro trial: *C*_2_ = −*β*_*c*_(*tanh*((*V*_*Pro-R*_ - *V*_*Pro-L*_)/*θ*_2_))^2^, and for an Anti trial: *C*_2_ = −*β*_*c*_(*tanh*((*V*_*Pro-L*_ - *V*_*Pro-R*_)/*θ*_2_))^2^.

*θ*_0_ is a fixed parameter that controls the sensitivity of this term, and *β*_*c*_ is a fixed parameter controlling the strength of this term. We used the fixed parameter values *θ*_1_ = 0.05, *θ*_2_ = 0.15, *β*_*c*_ = 0.001.

### Model Optimization

We initialized many different model solutions with random parameter values, and a random seed for the random number generator to generate unique noise for each model solution. For each initialization we minimized the cost function using constrained parabolic minimization.

#### Constrained Parabolic Minimization

The minimization starts by creating a local search radius, which restricts the scope of the search on each step. At each step, the algorithm approximates the cost function locally using the hessian matrix, and gradient vector, which defines a 16 dimensional parabolic surface. The minimization takes a step in the direction that minimizes the cost on this parabola subject to the constraint that the step length equals the search radius. If the resulting step would increase the cost function the step is not taken, the search radius is reduced, and another step is attempted. As the search radius becomes smaller, this method converges to gradient descent.

#### Two Stage Optimization

For each parameter initialization, an initial minimization was done using 50 trials/condition. If this initial minimization passed a set of criteria then a further minimization was done using 1000 trials/condition. The initial criteria was used to prevent long optimizations on model solutions that were performing very badly. The initial criteria was that performance on Pro trials was greater than Anti trials, and Anti performance on delay-period opto trials was worse than control or choice-period opto trials. The final minimization terminated after 1000 iterations, or when a step in parameter space reduced the cost function by less than 1e-12. When the minimization ended, if the final cost was below a threshold of −0.0001 we accepted the final parameter values as a model solution. We ended up with N=373 unique model solutions.

### Model analysis

To examine the space of model solutions, we clustered each model solution based on the dynamics of their simulated units. We simulated 200 trials for each model solution for each trial type (total 6 = Pro/Anti × control/delay-period opto/choice-period opto). Then, we computed the average trajectory for each unit in the model on correct and incorrect trials for each trial type. The average trajectories were concatenated into a model response vector (length *M* = 4 units × hit/miss × Pro/Anti × control/delay/choice opto × *T* timesteps). We created response matrix (*R*) of response vectors for all model solutions. (Size *N x M*). We used the Singular Value Decomposition to factor *R* = *U* * *S* * *V′*. The orthonormal matrices *U* and *V*′ are the directions of greatest variance in *R* across model solutions (*U*), and across time points (*V’*). The column space of *U* gives us the weights for each model solution onto the set of temporal basis vectors in the row space of *V’*. To compute the average task and choice decoding, we computed the d’ sensitivity index for each solution. We then computed the decoding accuracy via the formula: % correct = NormalCDF(d’/sqrt(2)).

## ACKNOWLEDGEMENTS

We thank K. Osorio and J. Teran for animal and laboratory support. This work was funded by Howard Hughes Medical Institute. C.A.D. was supported by a Howard Hughes Medical Institute predoctoral fellowship. C.A.D. and M.P. are currently supported by the Simons Collaboration on the Global Brain postdoctoral fellowship.

## AUTHOR CONTRIBUTIONS

C.A.D. collected electrophysiological and optogenetics data. M. P. and C.A.D. analyzed electrophysiological data. C.A.D. analyzed the optogenetics data. M.P., A.T.P., C.D.B. and A.J.R. generated and analyzed modeling results. A.A. and C.D.K. carried out the acute optogenetics experiments. J.C.E. and C.D.K. played an advisory role on electrophysiological and optogenetics experiments respectively. C.A.D., J.C.E. and C.D.B. conceived the project. C.A.D., M.P., A.T.P. and C.D.B. wrote the paper with comments from J.C.E.. C.D.B. was involved in all aspects of experimental design and data analysis.

---

**Figure S1.**
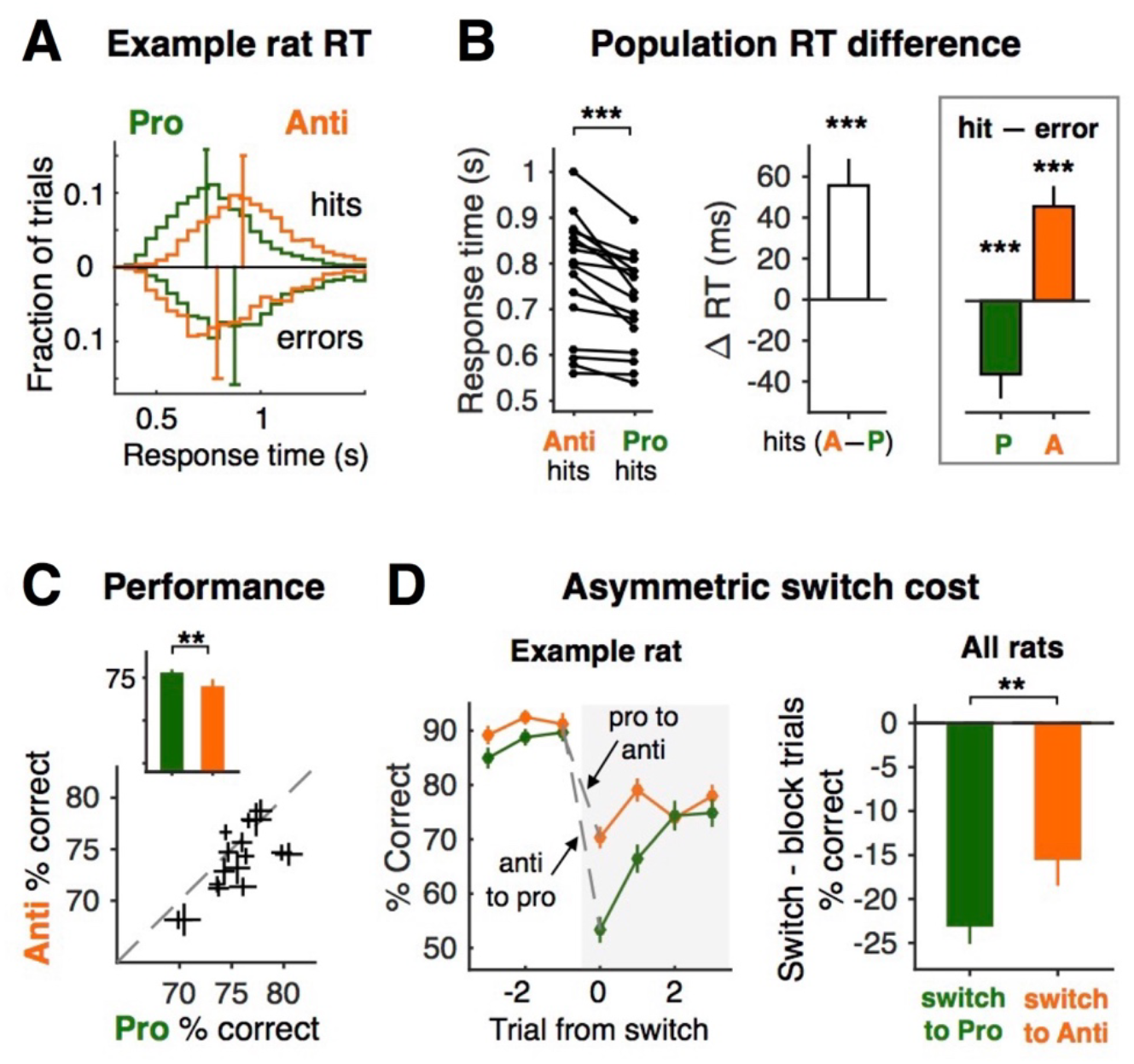
Post-surgery performance for implanted rats, Related to Figure 1. Asymmetries between Pro and Anti response time (RT), accuracy, and task switch cost observed in implanted rats replicate those found in freely moving rats in (Duan et al., 2015). (A) Normalized RT distributions of an example rat. Histograms of Pro and Anti RTs are shown here for hits (top) and errors (bottom). Each curve is normalized to have a total area of 1. Median RTs for Pro and Anti hits and errors are indicated by vertical bars; 95% confidence intervals across trials for each trial type are indicated by horizontal bars. (B) RT summary of 16 individual rats (7 for neural recordings and 9 for optogenetic inactivation experiments). Left: median RTs for Anti hits and Pro hits for all rats. Right: RT difference between Pro and Anti, hits and errors, averaged across all rats. For each rat, the difference between median RTs of paired conditions was calculated. White bar shows the mean and SEM across rats for Anti hit RTs minus Pro hit RTs. Green bar shows Pro hit RTs minus Pro error RTs. Orange bar shows Anti hit RTs minus Anti error RTs. (C) Pro and Anti performance for individual rats. Means and SEMs of Pro and Anti performance are computed over sessions for each rat and plotted against each other. Average Pro (green) and Anti (orange) performance across rats was plotted in the upper left corner. (D) Switch cost asymmetry. Left: percent correct as a function of trial number relative to a task block switch for one example rat. Each data point is the mean and SEM across trials for Pro and Anti accuracy on three trials before and after the switch. Right: average accuracy switch cost for Pro trials and Anti trials across rats. **p < 0.01; ***p < 0.001.

**Figure S2.**
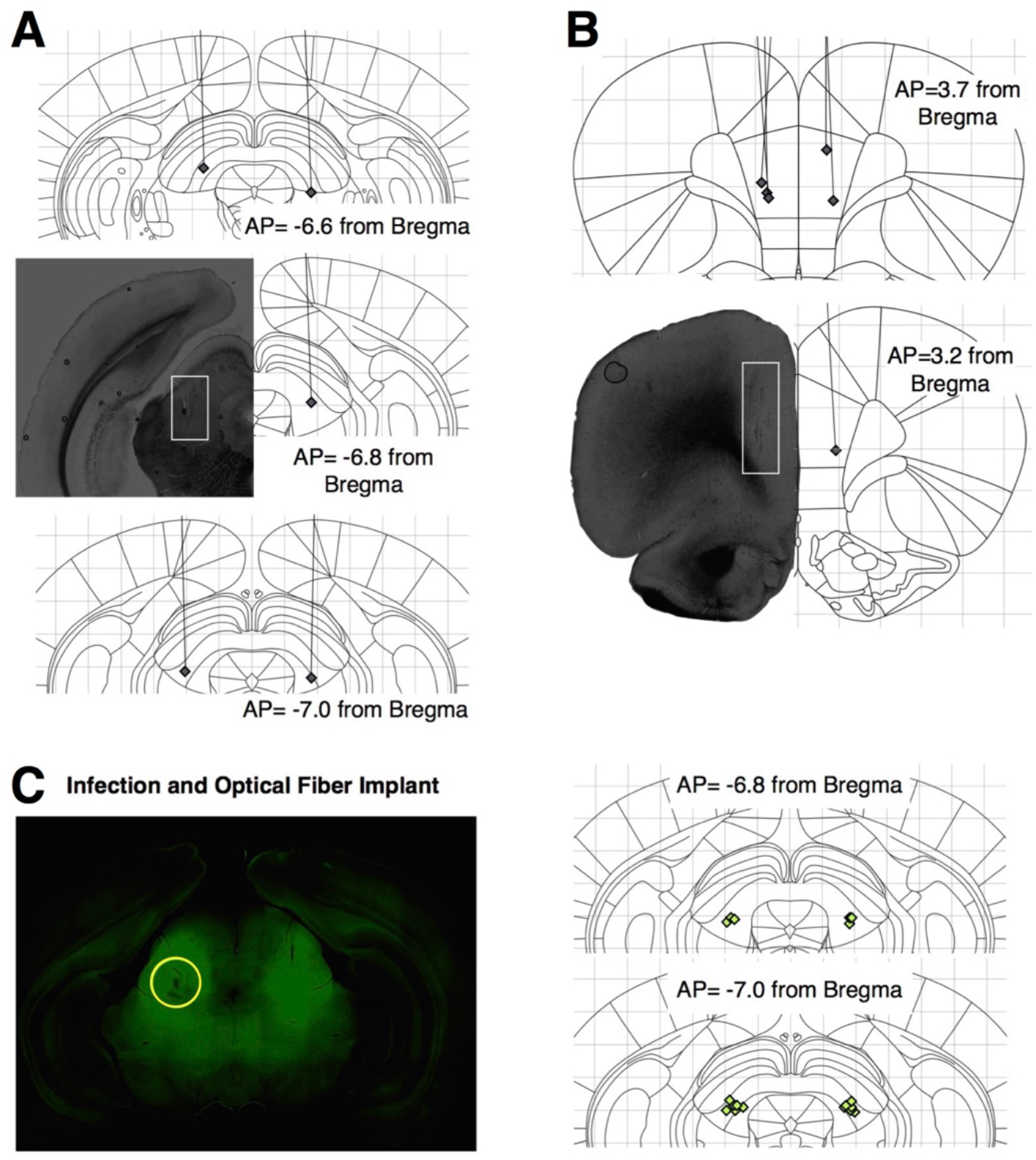
Histology for tetrode and optical fiber implantation, Related to Figure 1. (A) Histology for 5 rats with left or right SC tetrode implants. Gray diamonds indicate the final location of the tetrode tips. Lines indicate the tetrode tracks. (B) Histology for 6 rats with left or right mPFC tetrode implants, similar to A. Seven rats were implanted with tetrode drives all together: 4/7 rats with both SC and mPFC drives; 2/7 rat with only mPFC drives; 1/7 rat with only SC drive. (C) Histology for 9 rats with bilateral SC AAV virus infection and optical fiber implants. Left: example of AAV5-CaMKIIα-eYFP-eNpHR3.0 infection. Green fluorescence indicates the infection coverage. Yellow circle indicates the estimated spread of light stimulation based on previous acute recording experiments (Hanks et al., 2015). Right: green diamonds indicate the tips of etched optical fibers for all animals.

**Figure S3.**
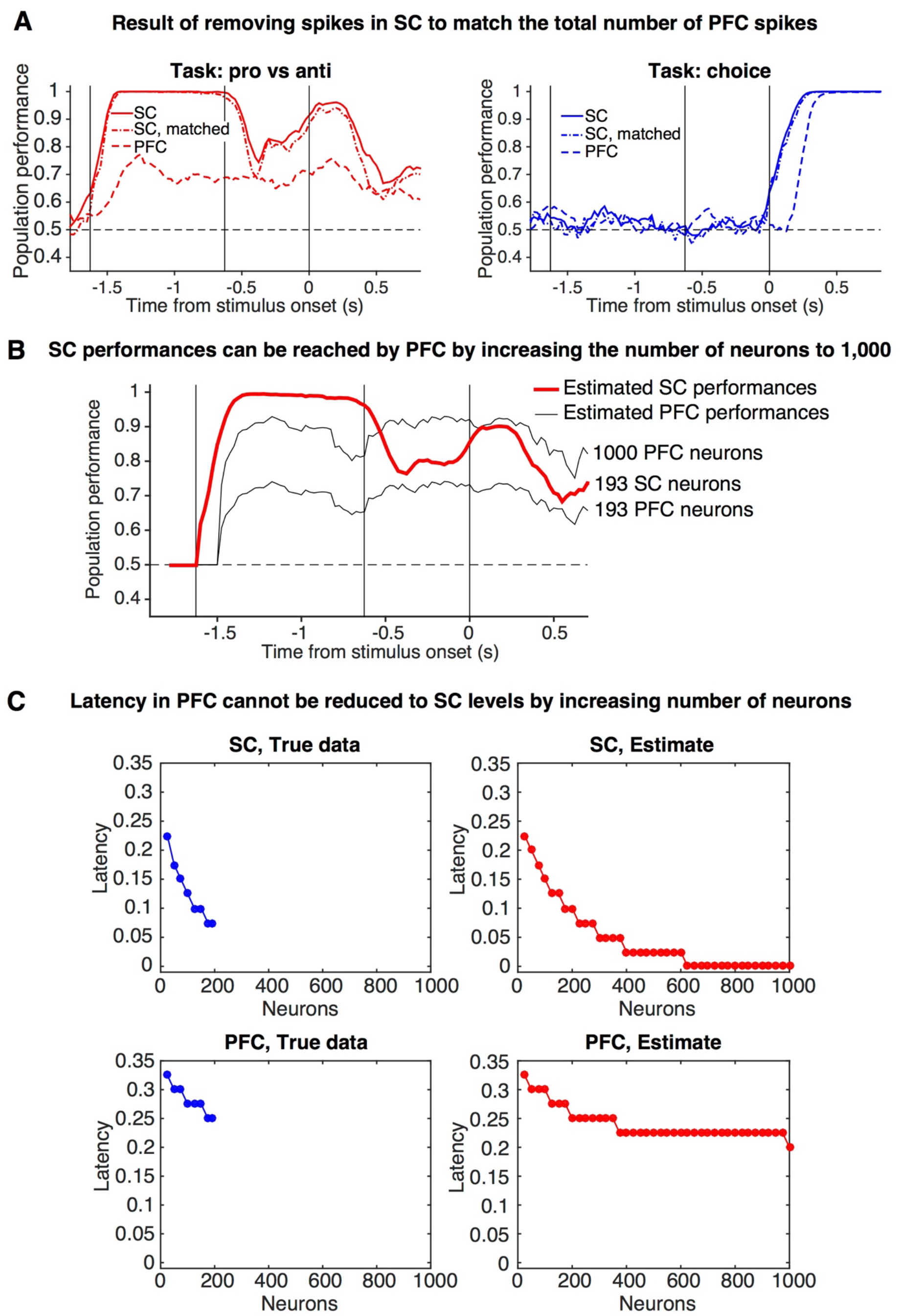
Controls for the comparison of population performances in SC and PFC, Related to Figure 2. (A) Pro/Anti (red, left) and Choice (blue, right) classification performances in SC (solid line), PFC (dashed line), and SC after matching the average firing rate by randomly removing spikes (dash-dot line). Pro/Anti performances in SC are still significantly higher than PFC after matching firing rates (p<0.05). Latency of the rise in choice classification is still shorter in SC after matching firing rates (p<0.01). (B) Estimated Pro/Anti classification performances for different number of neurons in SC (red) and PFC (black) (see STAR Methods). Performances of 193 SC neurons can be matched by approximately 1000 PFC neurons (C) Estimated latency of choice performances in PFC (blue, left) and in SC (red, right) for different numbers of neurons (see STAR Methods). SC latency cannot be matched by PFC even when considering a population of 1000 neurons.

**Figure S4.**
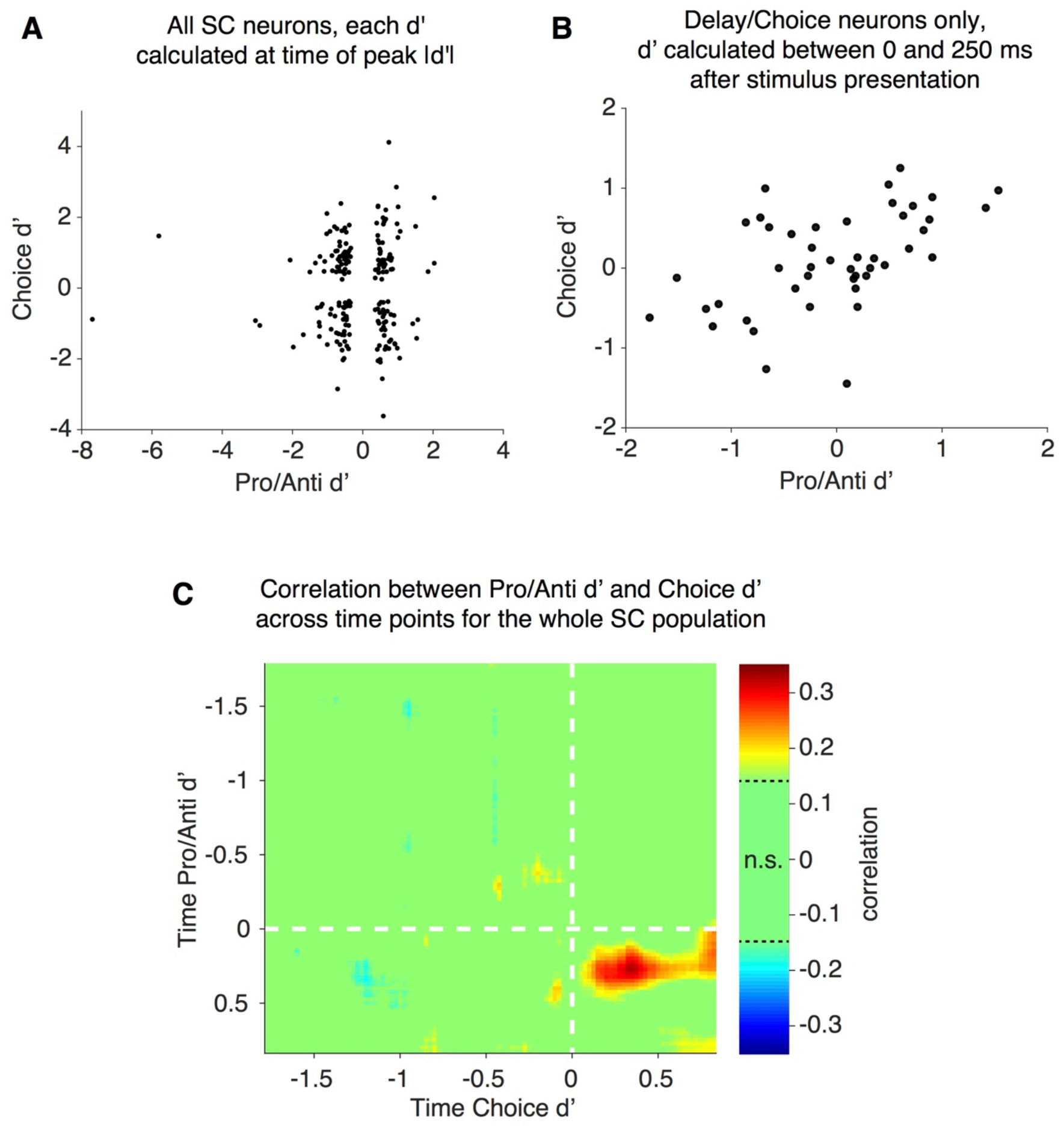
Relationship between task (Pro/Anti) and choice (Contra/Ipsi) d’ across the SC population, Related to Figure 4. (A) For each SC neuron, the signed Pro/Anti d’ computed at the time of peak Pro/Anti selectivity was plotted against the signed Choice d’ computed at the time of peak Choice selectivity. No correlation is observed (r = 0.06, n = 193, p>0.1). (B) For SC Delay/Choice neurons, the signed Pro/Anti d’ was plotted against the signed Choice d’, both computed within the first time bin after stimulus appearance (0-250 ms). The two are significantly correlated (r = 0.52, n = 45, p<10-4), due to the prevalence of Pro/Contra and Anti/Ipsi units. (C) Correlation between Pro/Anti d’ and Choice d’ for the whole SC population computed at all time points. The correlation is significantly different than 0 only at times shortly after the appearance of the target stimulus.

**Figure S5.**
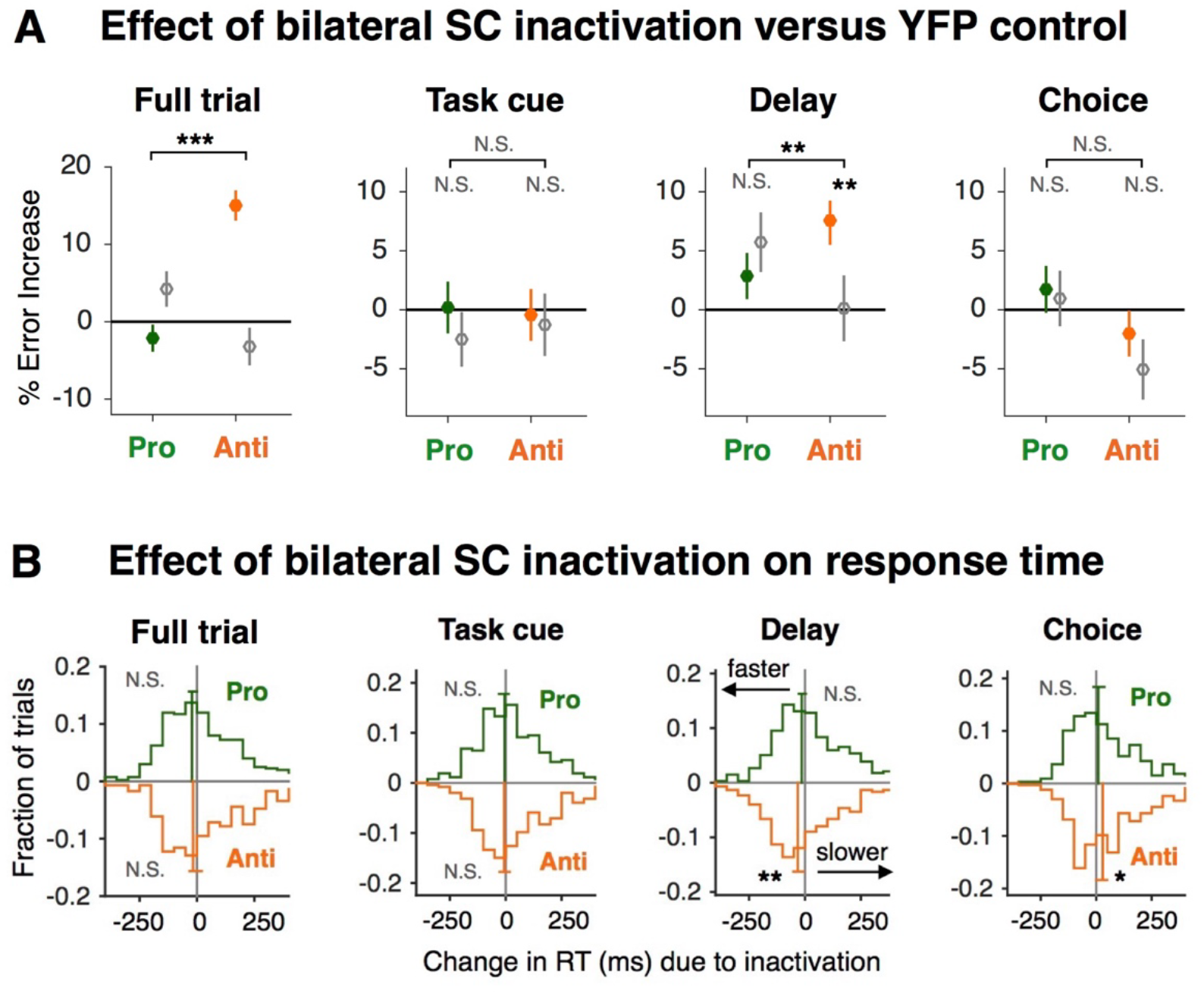
Effect of bilateral SC inactivation and YFP control, Related to Figure 5. (A) Effect of full-trial and sub-trial inactivations of bilateral SC on Pro (green) and Anti (orange) error rate (mean and s.e.m.) compared to YFP controls (gray). All paired statistics shown here are computed using a permutation test, shuffled 5000 times. (B) Effect of full-trial and sub-trial inactivations of bilateral SC on response time (RT). For each behavioral session, a median RT on non-stimulated control trials is calculated and subtracted from the RTs on inactivation trials, and these normalized RT changes due to inactivation are plotted here. Each curve is normalized to have a total area of 1. Vertical bars show the median RT changes for correct Pro and Anti trials; 95% confidence intervals across trials for each trial type are indicated by horizontal bars. A shift to the right indicates slowing due to inactivation and a shift to the left indicates speeding. N.S. p>0.05; *p<0.05; **p < 0.01; ***p < 0.001.

**Figure S6.**
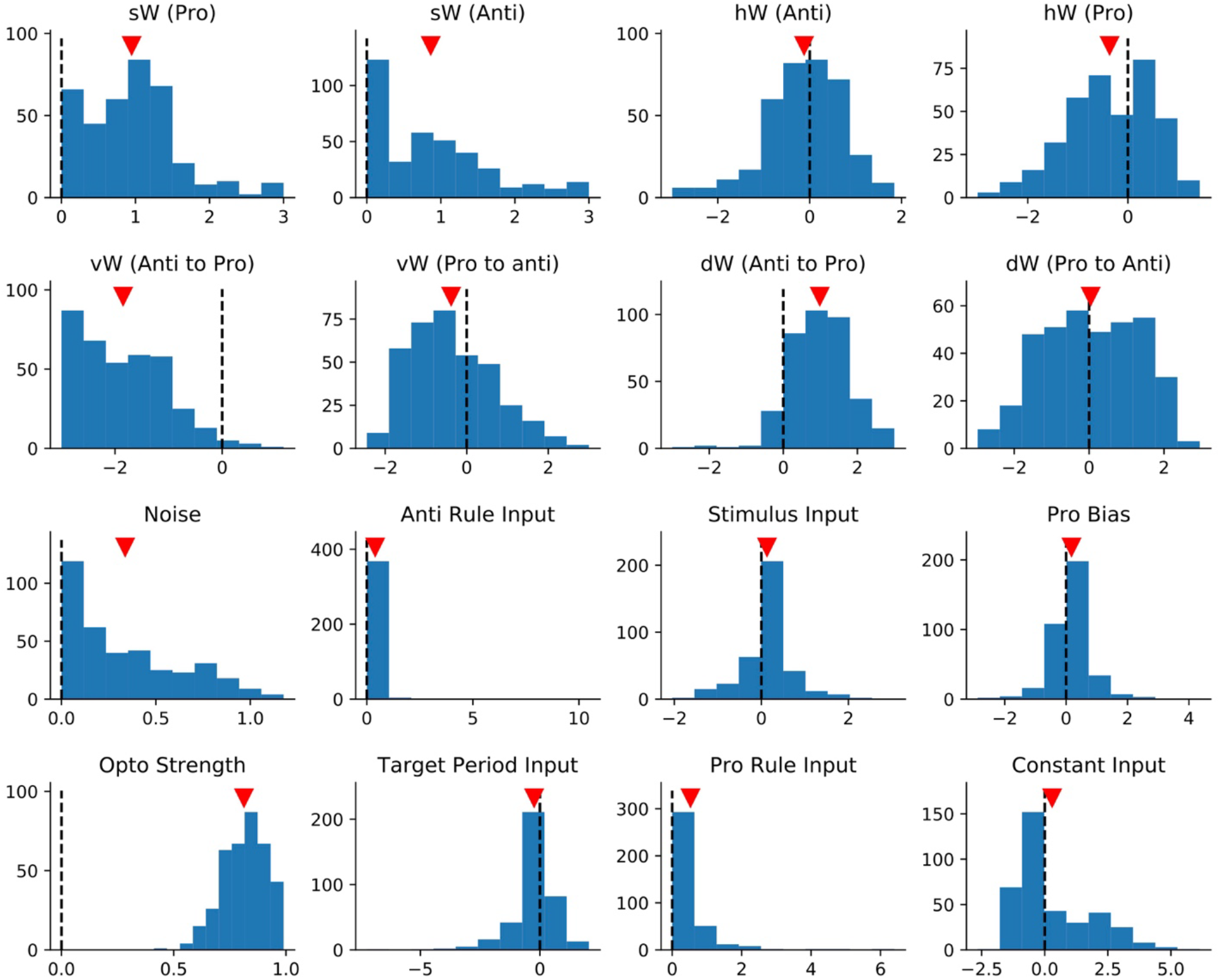
Distributions of parameters across all model solutions, Related to Figure 7. The distribution of parameter values across all solutions is plotted for each of the 16 free parameters. Vertical dashed line marks zero for reference. Red arrow marks average parameter value across solutions. The weight parameters determined the connectivity matrix between units. The noise parameter was the variance of white noise added to each unit on each time step. The Pro and Anti rule input weights determined the strength of the task inputs to either the Pro or Anti units. The stimulus input determined the weight of the stimulus to either the Left or Right units. The Pro bias term was a constant input to only the Pro units. The target period input was a constant input to all nodes, only during the target period. The constant input was a bias term during all time points for all units. The opto strength was the fraction of each node’s output that was transmitted to the other nodes during inactivations; a strength of 1 is no inactivation, a strength of 0 is complete inactivation.

**Figure S7.**
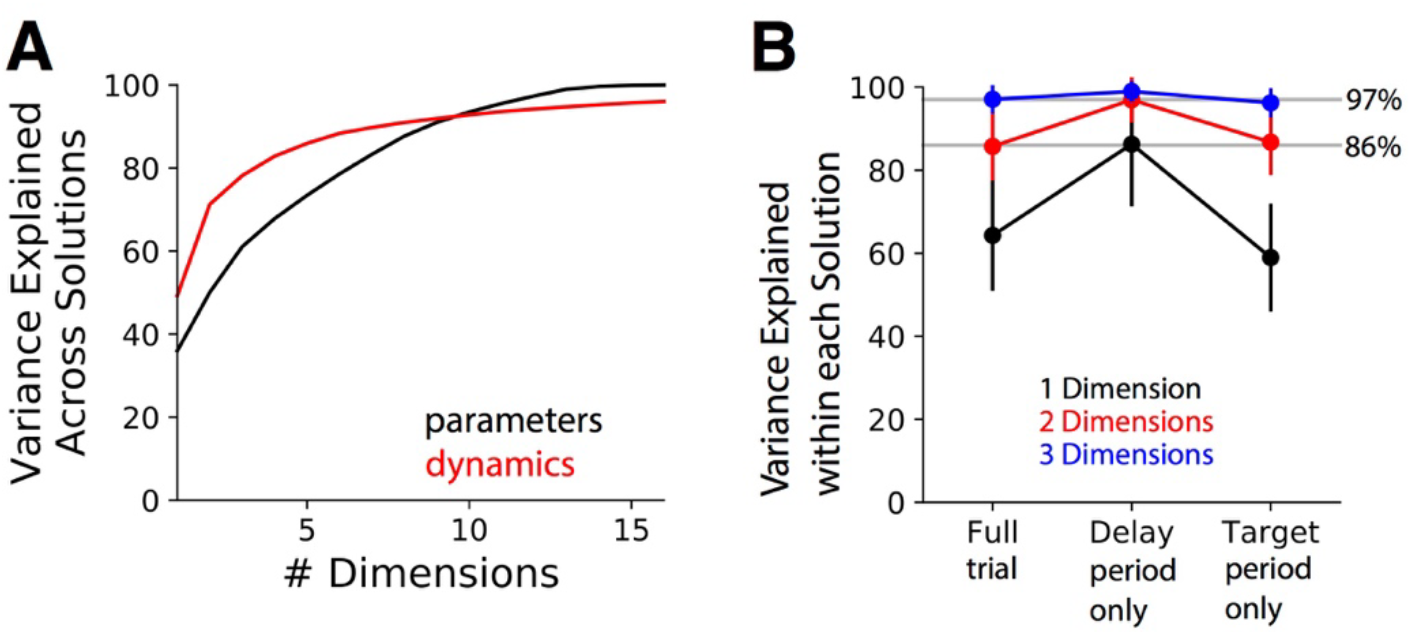
Variability across solutions in model dynamics and parameter space, Related to Figure 7. (A) The dimensionality of parameters across model solutions, and of dynamics across model solutions. 8 SVD dimensions are required to explain 90% of the variance in dynamics across model solutions. 10 PCA dimensions are required to explain 90% of the variance in parameters across model solutions. (B) Variance explained by each dimension of PCA performed on each trial’s dynamics. Full trial - PCA computed on all time points. Delay period only - PCA computed only during the delay period. Target period only-PCA computed only during the target period. Mean + s.d across model solutions.

